# From temperate to polar waters: Transition to non-cyanobacterial diazotrophy upon entering the Atlantic gateway of the Arctic Ocean

**DOI:** 10.1101/2025.01.27.635040

**Authors:** Lisa W. von Friesen, Christien P. Laber, Bjarke H. Kristensen, Emma Nysted, Marcus Sundbom, Stefan Bertilsson, Pauline Snoeijs-Leijonmalm, Hanna Farnelid, Lasse Riemann

**Affiliations:** University of Copenhagen, Department of Biology, Marine Biology Section, Strandpromenaden 5, DK-3000 Helsingør, Denmark; Linnaeus University, Centre for Ecology and Evolution in Microbial Model Systems (EEMiS), Kalmar, Sweden; Stockholm University, Department of Environmental Science (ACES), Stockholm, Sweden; Swedish University of Agricultural Sciences, Department of Aquatic Sciences and Assessment, Uppsala, Sweden; Stockholm University, Department of Ecology, Environment and Plant Sciences, Stockholm, Sweden

## Abstract

Nitrogen fixation, the microbial reduction of dinitrogen to ammonia, is increasingly recognized to occur in the Arctic Ocean. However, knowledge about the composition, biogeography, abundance, and ecology of nitrogen-fixing organisms (diazotrophs) is poor. This ultimately hinders prediction of ecosystem productivity fueled by nitrogen fixation in this rapidly changing and predominantly nitrogen-limited ocean. We assessed the composition and abundance of total and active diazotrophs in sub-surface water (8 m; amplicon sequencing and quantification of the marker gene *nifH*) over ∼3,400 km from the mouth of the brackish Baltic Sea to the sea ice edge in the Arctic Ocean. Upon entering nutrient-rich waters in the Atlantic gateway to the Arctic, we discovered an abrupt transition from autotrophic to heterotrophic diazotrophic activity. Our findings suggest that diazotrophy is functionally distinct in the Arctic Ocean compared to adjacent temperate-boreal waters – a difference likely driven by inorganic nutrients, salinity, and temperature. We identify three key non-cyanobacterial diazotroph groups in the Arctic Ocean with Arctic-specific (Rhodocyclales and Oceanospirillales) or more widespread (unknown Gammaproteobacterium) distribution patterns and report their *nifH* gene transcription levels (up to 10^3^ *nifH* transcripts L^−1^). In contrast, actively transcribing diazotrophs in the warmer and more nutrient-poor Norwegian Sea with coastal-influenced water were dominated by sublineages of Candidatus *Atelocyanobacterium thalassa* (UCYN-A1, UCYN-A2, UCYN-A4; up to 10^4^ *nifH* transcripts L-1). With ongoing atlantification of the Arctic pushing oceanic provinces and biogeographical ranges poleward, we predict a future displacement of the transition from autotrophic to heterotrophic diazotrophic activity with likely significant changes in nitrogen fixation.

## Introduction

Nitrogen fixation is the biological reduction of molecular nitrogen to ammonia by a diverse functional group of microorganisms called diazotrophs. Cyanobacterial diazotrophs produce organic carbon and acquire energy through oxygenic photosynthesis (photoautotrophic), whereas, in contrast, the poorly understood non-cyanobacterial diazotrophs (NCDs) are thought to mostly utilize organic substrates and/or alternative energy and carbon sources (chemoheterotrophic) (Turk-Kubo et al. 2022). By stimulating new primary production through the supply of nitrogen, diazotrophs enhance carbon sequestration (Karl et al. 1997) and are thereby key players in the biogeochemical cycling of nitrogen and carbon in the global ocean (Zehr and Capone 2020).

Evidence of nitrogen fixation in the Arctic Ocean is accumulating with observations of diazotrophic cyanobacteria and NCDs, but their activity, biogeography, abundance, and environmental controls are largely unknown (von Friesen and Riemann 2020). This hampers our ability to predict and understand what governs nitrogen availability and, in turn, influences primary production potential in the Arctic Ocean. There are indications of distinct community patterns, unique phylotypes, and size distributions among Arctic diazotrophs (Farnelid et al. 2011; Fernández-Méndez et al. 2016; Salazar et al. 2019; Pierella Karlusich et al. 2021; Shiozaki et al. 2023) – similar to distinctions between polar and non-polar microbial communities (e.g. Ghiglione et al. 2012). These biogeographical patterns are likely governed by the distinctive environmental characteristics of the Arctic Ocean – a region influenced by sea ice, riverine and glacial input, low temperatures, and strong seasonality.

The Arctic Ocean is strongly impacted by inflow from the Atlantic and Pacific Oceans, creating sharp environmental gradients that are well suited for deciphering drivers of diazotroph community composition and activity transitions. For instance, in a transect sampling from the equatorial Pacific Ocean through the Bering Strait into the Arctic Ocean, most diazotrophs gradually disappeared northwards, except for the diazotroph Candidatus *Atelocyanobacterium thalassa* (UCYN-A) (Shiozaki et al. 2017). UCYN-A is widespread and contributes significantly to global nitrogen fixation (Farnelid et al. 2016) and was until recently thought to be a photoheterotrophic cyanobacterial symbiont with a haptophyte host but is now established as an early stage of a eukaryotic organelle (the nitroplast; Coale et al. 2024). Due to its association with a phytoplankton cell, we here refer to UCYN-A as a photoautotrophic-associated diazotroph, although the diazotroph (or organelle) in itself does not fix carbon. In the Pacific Arctic, UCYN-A fixes nitrogen at rates similar to those reported from non-polar regions (Harding et al. 2018). However, once departing from the Pacific Arctic shelves and entering the high Arctic, putative anaerobic NCD phylotypes had high relative abundances also at locations where nitrogen fixation rates were measured and UCYN-A was absent (Blais et al. 2012; Shiozaki et al. 2018). In the Canadian Arctic gateway outflow, NCDs dominated and exhibited distinct biogeography (Robicheau et al. 2023b), similar to patterns described at coastal locations in Svalbard and Greenland (von Friesen et al. 2023). Hence, the following hypotheses arise: 1) NCDs are significant contributors to nitrogen fixation in Arctic regions, and 2) diazotrophs of the Arctic Ocean are distinct from those in adjacent seas. Currently, data on a south-north transition in the distribution of cyanobacterial diazotrophs and NCDs does not exist. Due to their distinct functionality (i.e. phototrophy versus heterotrophy), knowledge of the distribution and regulation of diazotrophs is critical for understanding current and predicting future nitrogen fixation in the Arctic Ocean.

Compared to the Pacific inflow, knowledge of diazotroph composition and activity (diazotrophy) in the Atlantic inflow is scarce. The Atlantic inflow is about 10 times larger than the Pacific inflow and is ecologically important with observed increasing inflow of Atlantic water to the Arctic Ocean (“atlantification”), advecting warm, saline, and nutrient-rich waters and temperate-boreal species northwards (Årthun et al. 2012). Intensified atlantification is expected to stimulate primary production in the Arctic Ocean, but the availability of nitrogen to fuel such a continued increase is uncertain (e.g. Lewis et al. 2020). Since nitrogen fixation is a source of new nitrogen to the ocean, understanding the environmental drivers governing diazotrophy along the Atlantic inflow to the Arctic is a research priority.

We analyzed the diazotroph abundance (quantitative PCR of *nifH,* a functional marker gene for diazotrophs, from DNA and cDNA), composition, and transcriptional activity (*nifH* amplicon sequencing of DNA and cDNA) in sub-surface water (∼8 m depth) over a ∼3,400 km long transect covering the Atlantic inflow to the Arctic Ocean. The transect sampling was conducted in two seasons (summer and autumn), traversing contrasting environmental conditions across the North Sea, Norwegian Sea, and Barents Sea: from the mouth of the temperate-boreal Baltic Sea in the south to the marginal sea ice zone of the Arctic Ocean in the north. We expected latitudinal differentiation in diazotroph community composition, abundance, and *nifH* transcription, with an increasing proportion of NCDs relative to cyanobacteria towards polar waters. We aimed to assess underlying environmental and ecological drivers by coupling their biogeography, transcription, and abundances with prokaryotic community composition (16S rRNA gene amplicon sequencing) and physicochemical and biological variables (inorganic nutrients, temperature, salinity, fluorescence, and cell abundances).

## Materials and

### Study region and sampling

The objective was to study diazotroph community structure and activity along a transect reaching between temperate-boreal to polar waters. The region is strongly influenced by the North Atlantic Current originating from the Gulf Stream and transitioning northward along and into the Norwegian Coastal Current, where it mixes with coastal water (Fig. 1A). Further north, it branches either eastward over the Barents Sea where it meets the cold Bear Island and East Spitsbergen Arctic currents or into the West Spitsbergen Current reaching around the north-west corner of Svalbard into the Arctic Ocean to the east (Fig. 1A).

**Fig. 1.**
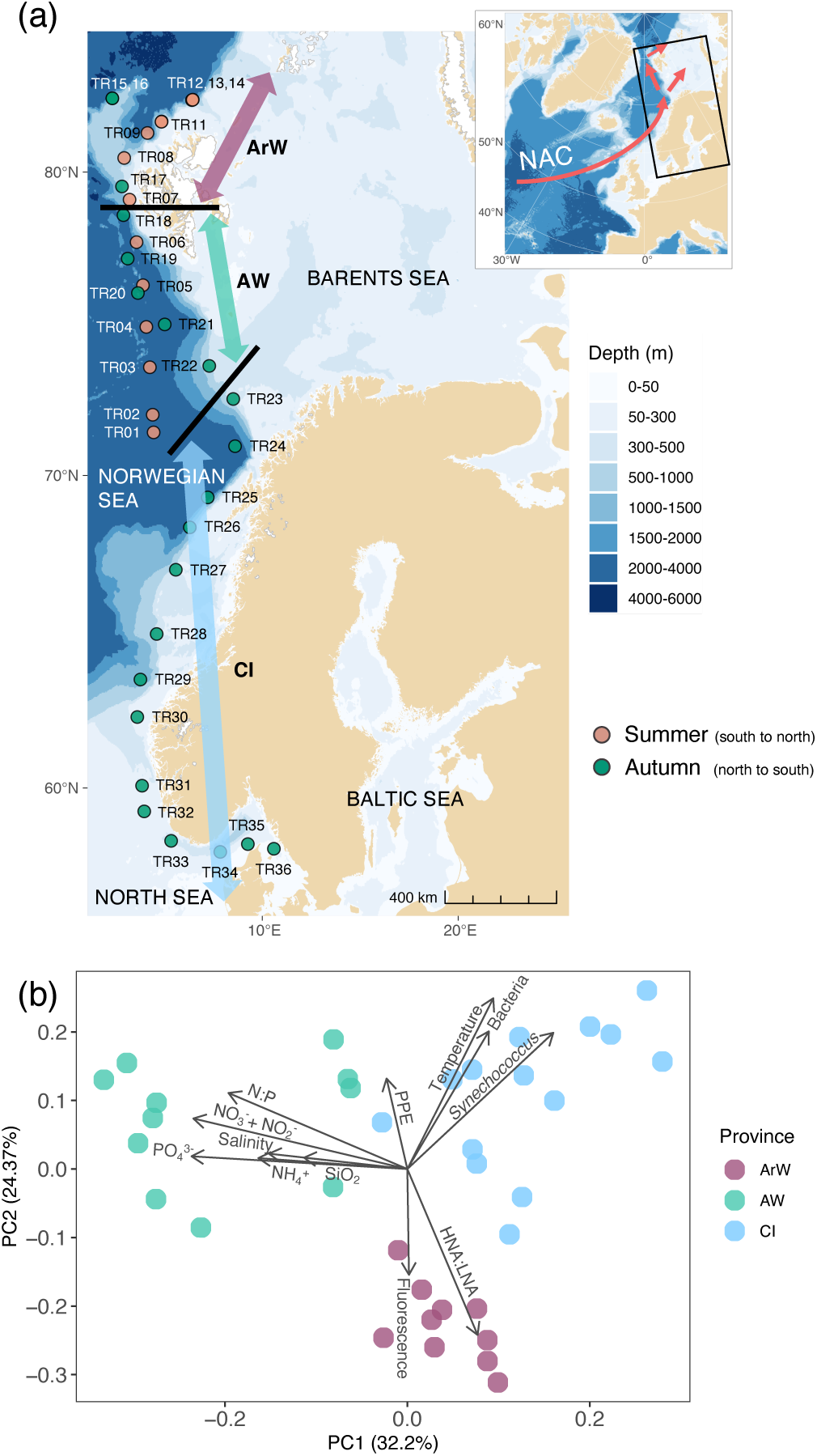
Map of the study area and definition of ecological water provinces. ArW: Arctic Water, AW: Atlantic Water, CI: Coastal Influenced Water. **(a)** Sampling stations (TR) in summer (coral: south to north) and autumn (green: north to south). Black thick lines denote the division between provinces. The insert shows the study region on a more zoomed-out map of the North Atlantic. NAC: North Atlantic Current, NCC: Norwegian Coastal Current, WSC: West Spitsbergen Current. The red arrows depict warmer Atlantic currents and the blue arrow colder Arctic currents. The bathymetry legend is not valid for the insert. **(b)** Principal Component Analysis (PCA) of environmental metadata for each station consolidating the definition of ecological water provinces. Displayed as the two first principal components. PO_4_^3-^: phosphate, NO_3_^−^ + NO_2_^−^: nitrate + nitrite, NH_4_^+^: ammonium, SiO_2_: silicate, PPE: photosynthetic picoeukaryote cells ml^−1^, HNA:LNA: the ratio between high nucleic acid and low nucleic acid bacterial cells (a proxy for bacterial activity), N:P: the molar ratio between dissolved inorganic nitrogen and phosphorus.

Samples were collected during the Synoptic Arctic Survey (IB Oden, 2021; Supporting Information Table S1; Data S1; Swedish Polar Research Secretariat, 2022). All station names start with “TR”, denoting “transect”. Summer sampling (29 July – 2 August; stations TR01-TR14; south to north) ranged from the northern Norwegian Sea to north of Svalbard, and autumn sampling (11 – 18 September; stations TR15-TR36; north to south) ranged from north of Svalbard to the northern Kattegat (mouth of the Baltic Sea) (Fig. 1A). Sub-surface seawater (∼8 m) was obtained through the vessel’s continuously running underway water-intake system. Sampling time was adjusted relative to daylight for each latitude – during autumn using nautical almanack tabulations targeting mid-day and 30 min before sunrise and sunset, while summer samples were collected every eight hours (∼05, 13, 21 UTC) because of the continuous daylight. Each sample collection event for nucleic acid extraction, cell enumeration, and inorganic nutrient measurements lasted approximately five minutes.

### Physicochemical measurements

Temperature and salinity (SBE-45, Sea-Bird Scientific, WA, USA) and fluorescence (a proxy for chlorophyll *a*; FLNTURT-1899, 470/695 nm, WET Labs, Sea-Bird Scientific) were continuously measured. Samples for PO_4_^3-^, NO_3_^−^ + NO_2_^−^, NH_4_^+^, and SiO_2_ were filtered in duplicate (0.22 μm, Sterivex, Millipore, MA, USA) and refrigerated until on-board 4-channel continuous segmented flow analysis with photometric detection (QuAAtro39, SEAL Analytical) with a handling time of less than a day.

### Cell enumeration by flow cytometry

Seawater from the start and end of each 5 min sampling event was pooled and fixed in 0.5% glutaraldehyde (Sigma Aldrich, MA, USA) combined with 0.01% Pluronic F-68 (Gibco, MA, USA) for picocyanobacterial and eukaryotic cells (Marie et al. 2014). Samples were incubated for 5 min at room temperature, frozen at -80°C, and autotrophic and heterotrophic cells enumerated (Alegria Zufia et al. 2021). Photosynthetic picoeukaryotes (PPE) were quantified using forward scatter as a proxy for cell size and red fluorescence as a proxy for chlorophyll-*a* content. *Synechococcus* were distinguished using the orange fluorescence signal as a proxy for phycoerythrin content (Cube8, Sysmex, UK; 488 nm laser). Bacteria were stained with 0.06% SYBR I (Invitrogen, MA, USA), and HNA (high-nucleic acid cell) and LNA (low-nucleic acid cells) cells were identified using side scatter and green fluorescence signals (CytoFlex, Beckman Coulter, USA; 488 nm laser) (Gasol and Morán 2015). Population gating was performed in FCSalyzer (v.0.9.22).

### Sampling and extraction of nucleic acids

Cells for DNA and RNA extractions were concentrated by pressure filtration (30 min; ∼1500 ml; 200 μm pre-filtration; 0.22 μm Sterivex). Triplicate filtrations were carried out in dim light, subsequently preserved in RNAlater (Ambion, CA, USA), and frozen at -80°C. Duplicate blank filters served as contamination controls covering the full process from sampling to sequencing. Additional water from Niskin bottles of a parallel CTD cast (10 m; TR15/16) (Fig. 1A) was collected to account for any potential microbial contamination from the underway water intake system. The prokaryotic and diazotroph community composition in samples obtained via the underway and with Niskin bottles were similar (16S rRNA gene: average Euclidean distance between control samples 43.5 as compared to an average distance to all other samples of 74.3; Fig. 2A, B) (*nifH* gene: average Euclidean distance between control samples 13.9 as compared to average distance to all other samples 21.3, Fig. 3A, B). All the top 50 *nifH* ASVs observed in the three underway replicate samples (TR16) were also detected in the CTD control sample (TR15) with similar community composition (>95% Gammaproteobacteria) (Fig. 3A, B). Thus, we conclude that there were no significant differences between the underway and the CTD samples.

**Fig. 2.**
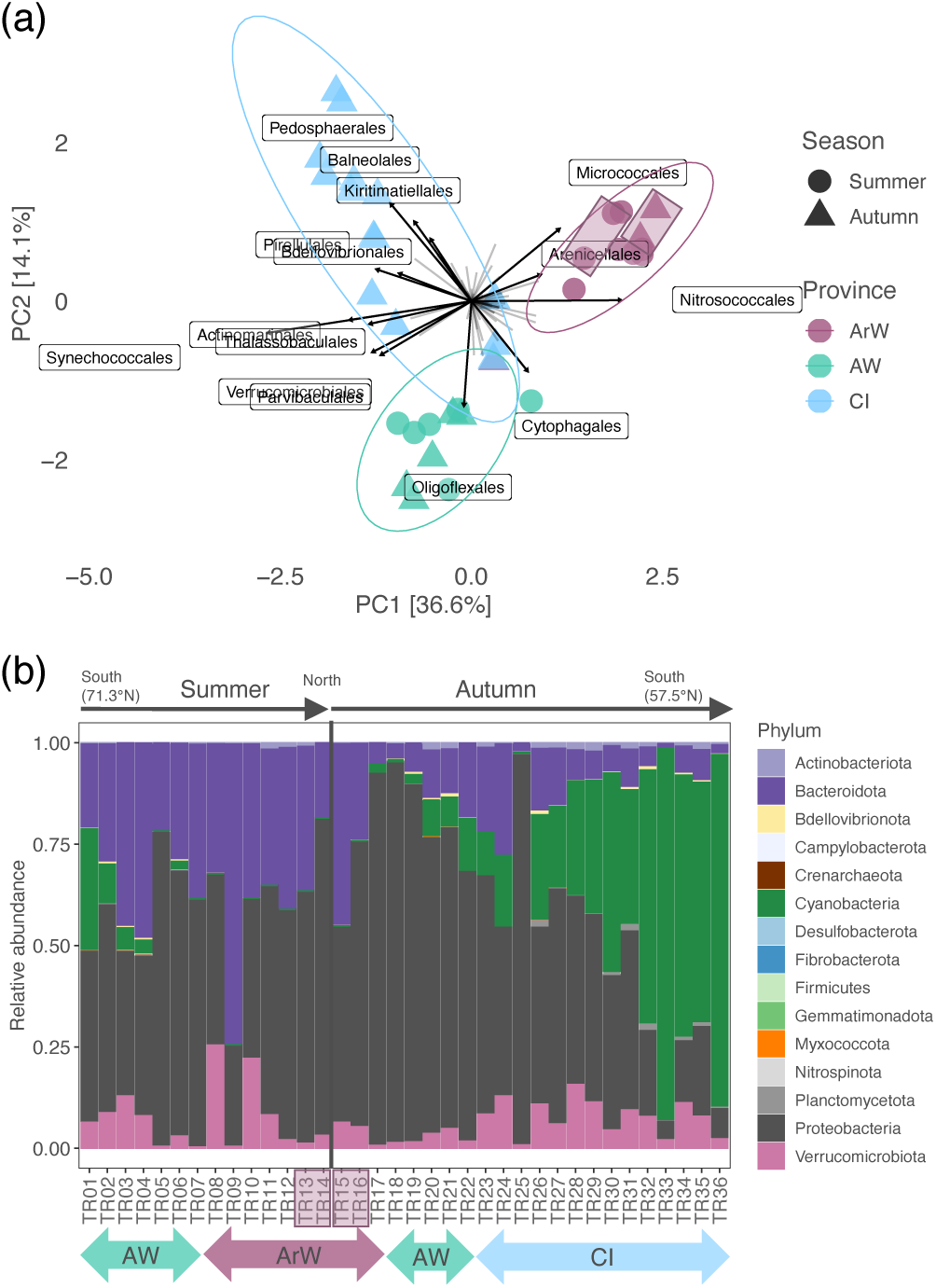
Prokaryotic community structure. Based on 16S rRNA gene amplicon sequencing along the summer and autumn transects. **(a)** Principal Component Analysis (PCA) on the order level displaying the top 15 influencing orders and their association with the three provinces. Light grey lines from the center represent the loadings of remaining orders (i.e. beyond the top 15). **(b)** The relative abundance of phyla along the transects. ArW: Arctic Water, AW: Atlantic Water, CI: Coastal Influenced Water. Note that CI was only sampled in autumn. Semi-transparent purple (ArW) boxes in A and B denote control sample pairs (CTD and underway water intake system in parallel).

**Fig. 3.**
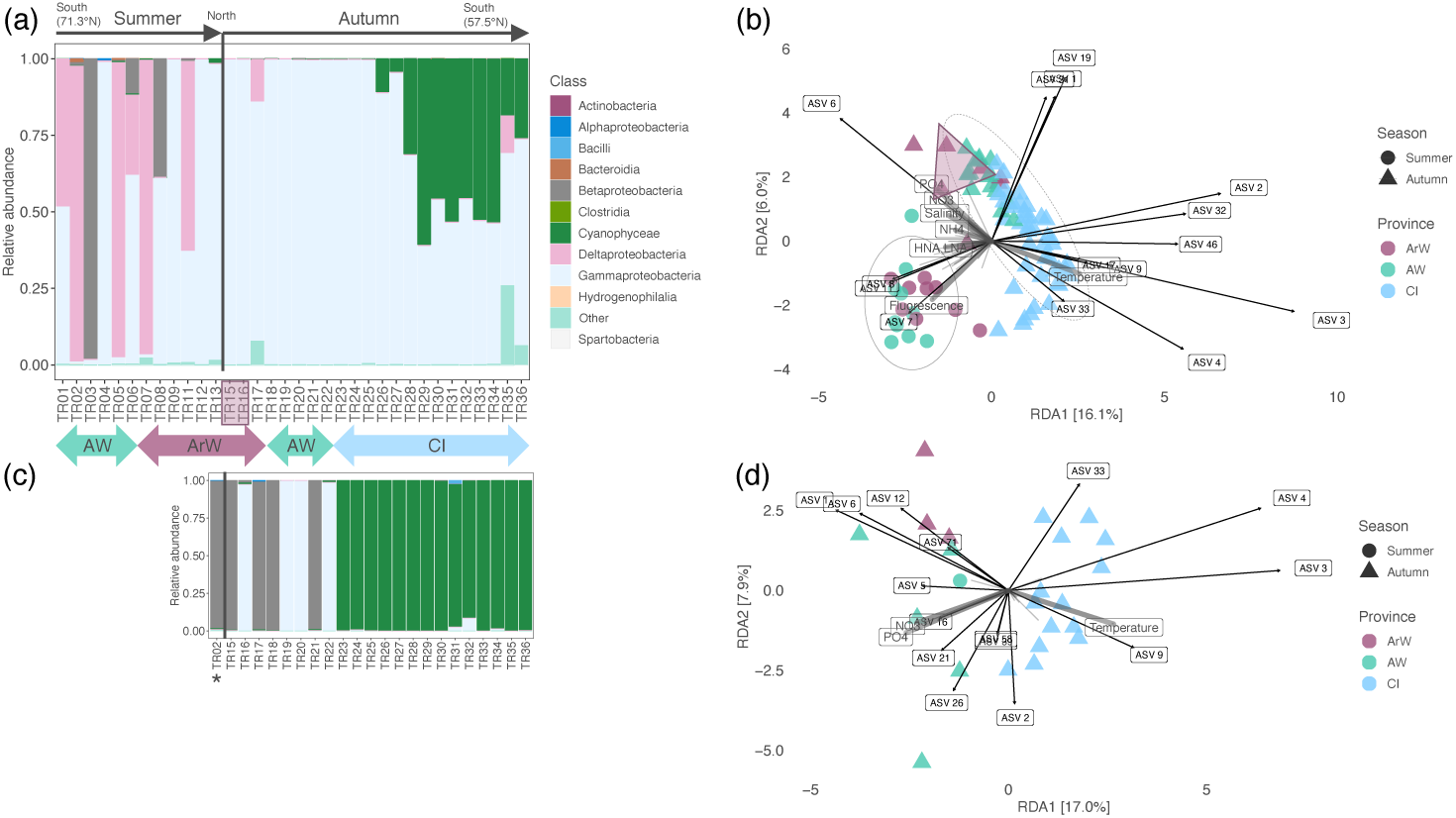
Diazotroph community structure. Composition of present (DNA) and actively transcribing (cDNA) diazotrophs based on *nifH* gene amplicon sequencing. **(a)** The relative abundance of classes along the summer and autumn transects (one to three replicates merged for each station). **(b)** Redundancy Analysis (RDA) on the amplicon sequence variant (ASV) level (top 50) displaying the top 15 influencing ASVs (black) and the significant environmental variables constraining the RDA (grey arrows). **(c)** The relative abundance of classes transcribed along the autumn transect. The asterisk denotes sample TR02 from summer AW. **(d)** RDA on the ASV level (top 50) displaying the top 15 influencing ASVs (black) and the significant environmental variables constraining the RDA (grey arrows). Refer to Fig. 4 for phylogenetic information on each ASV. ArW: Arctic Water, AW: Atlantic Water, CI: Coastal Influenced Water. Note that CI was only sampled in autumn. cDNA was only successfully sequenced from the autumn transect (all provinces), except for station TR02 from summer (*). PO_4_^3-^: phosphate, NO_3_^−^: nitrate + nitrite, NH_4_^+^: ammonium, HNA:LNA: the ratio between high nucleic acid and low nucleic acid bacterial cells (a proxy for bacterial activity). The semi-transparent purple box in a and the triangle in b denote control sample pairs (CTD and underway water intake system in parallel). The light grey lines from the center in b and d represent the loadings of remaining ASVs (i.e. beyond the top 15).

Before extraction, RNAlater was pushed out of the Sterivex filter, the casing removed, and the filter excised and subjected to five rapid freeze-thaw cycles with liquid nitrogen and thorough grinding with a sterile metal pestle. DNA and RNA were retrieved from combined extractions of the same filter (AllPrep DNA/RNA Mini kit, Qiagen Sciences, MD, USA) following the manufacturer’s instructions and quantified with PicoGreen and RiboGreen (Quant-iT, Invitrogen), respectively.

### 16S rRNA gene amplification and sequencing

16S rRNA gene fragments were amplified from DNA with primers 515F-Y and 926R (Supporting Information Table S2; 0.5 μM) targeting the V4 and V5 hypervariable regions using Phusion High-Fidelity PCR Master Mix (Thermo Scientific, MA, USA), indexed with Nextera DNA Dual-index N7-N5, magnetic bead-purified (AMPure, Beckman Coulter, CA, USA) and sequenced with Illumina MiSeqV3 (2×300 bp; National Genomics Infrastructure, Uppsala, Sweden). Sequences were quality controlled (Nextflow v.22.04.0; nf-core/ampliseq; v.2.3.0; Straub et al. 2020), assigned into ASVs (DADA2; v.1.20.0; Callahan et al. 2016) and taxonomically annotated (SILVA; v.132; Quast et al. 2013). Identification of UCYN-A and *Nodularia sp.* was enabled by blastn against known diazotroph 16S rRNA gene sequences compiled from GenBank (100% identity).

### cDNA synthesis

Synthesis of cDNA was performed with SuperScript IV (Invitrogen) in a two-step 10 μl reaction, with the first containing 5-5.5 μl undiluted RNA extract (10-1246 ng, median 82 ng), primer nifH3 (0.2 μM), MgCl_2_ (5 mM), and dNTPs (0.5 mM) at 65°C-300 s, 4°C-120 s. For the second step, SSIV buffer (1x) was added along with DTT (5 mM), ribonuclease inhibitor (2 U μl^−1^), bovine serum albumin (0.5 μg μl^.1^) and reverse transcriptase (10 U μl^−1^) at 50°C-50 min, 80°C-10 min.

### *nifH* gene amplification and sequencing

*nifH* amplicons were generated with nested PCR (Zehr and Turner 2001) using the primers nifH1-nifH4 (Supporting Information Table S2) and indexed as described elsewhere (von Friesen et al. 2023). The amount of template in the first stage (outer) PCR was as median 3.5 ng (0.5-42 ng) DNA or 1-5 μl cDNA. For each reaction, 1 μl (DNA) or 2 μl (cDNA) PCR product was then transferred to the second stage (inner) PCR. Each RNA sample was controlled for DNA contamination by using the equivalent amount of RNA extract as the template in the nested PCR, generating no visible amplification and no more reads than sequenced negative controls. Negative controls (PCR grade water) were included in each PCR run, and although with no visible amplification, five control samples were sequenced – facilitating a robust analysis of potential background contamination (see next section). Sequencing was performed with Illumina MiSeqV3 (2×300 bp; University of Copenhagen, Geogenetics, Denmark).

### Analysis of *nifH* amplicon sequence variants

Generation, quality control, and taxonomic assignment of amplicon sequence variants (ASVs) were performed as previously described (von Friesen et al. 2023). In brief: 1) DADA2 was used for ASV inference (trimming: 230 forward, 170 reverse), 2) selected steps of the NifMAP pipeline were applied to eliminate potential *nifH* homologs and translate sequences into amino acids (v.1.0; Angel et al. 2018), 3) assigning phylogenetic *nifH* clusters (Frank et al. 2016), and 4) taxonomic assignment against a *nifH* database (v.1.1.0; Moynihan 2020). A total of 936 ASVs were obtained with 421 - 136,249 (median 60,833) and 2 - 7,313 (median 984) reads per sample for environmental samples and controls, respectively. The R-package decontam (prevalence method; v.1.12.0; Davis et al. 2018) was applied, resulting in the identification and removal of four ASVs assigned as probable contaminants based on their prevalence in sequenced negative controls. ASVs remaining in negative controls were assigned to mostly (76%) unique ASVs not identified in environmental samples, and the remaining 24% corresponded to a median of 0.2% of the total read count in environmental samples. Samples with <1,000 reads (two samples) were removed after inspection of rarefaction curves as they did not reach an asymptote. Overall, *nifH* genes from between one and three replicate samples were successfully amplified, often exhibiting only small variations between each other. There were, however, occasional outliers, suggesting a patchy distribution of diazotrophs and/or heterogeneous water masses (Supporting Information Fig. S1A-B; median, min, and max distance to the median of multivariate dispersion was 16.9, 8.6, and 26.6, respectively). These results underline the importance of replicated sampling to obtain a representative description of the local diazotroph community.

The top 50 ASVs (91.5% of total reads) were queried against NCBI non-redundant databases (https://blast.ncbi.nlm.nih.gov/) to identify similar sequences, a maximum-likelihood amino acid phylogenetic tree was built in raxmlGUI (v.2.0.10; Edler et al. 2021; L-INS-I alignment; LG; G4 gamma mean; 100 bootstraps) and visualized with iTOL (https://itol.embl.de; Letunic and Bork, 2021). In addition, sequences from a recent NCD database were included in this tree (Turk-Kubo et al. 2022). The phylogeny of UCYN-A sublineages was assessed through nucleotide alignment and phylogenetic tree construction of Chroococcales-assigned ASVs (among the top 100 ASVs) with UCYN-A reference sequences (Farnelid et al. 2016; Turk-Kubo et al. 2017) in GeneiousPrime (v.1.1; UPGMA tree; Jukes-Cantor; 1000 bootstraps) (Supporting Information Fig. S2).

The *nifH* gene has been questioned as a sole marker for diazotrophs (Mise et al. 2021). However, uncertainties revolve mainly around Clostridia, which in our study represented only 0.0006% of total relative *nifH* read abundances. We judge this as negligible and see *nifH* as a reliable proxy for nitrogen fixation potential in our samples.

### qPCR and RT-qPCR of *nifH*

*nifH* gene- and transcript abundances were measured for select taxonomic groups using quantitative PCR (qPCR) and reverse transcription (RT)-qPCR (Supporting Information Table S3). Three new primer/probe sets were designed (MEGA X; v.10.1.7; Primer3Plus), targeting the ten *nifH*-transcribing NCD ASVs (grouped into, and from now on referred to as Beta-Arctic1, Gamma-Arctic1, Gamma-Arctic2) in the Atlantic gateway to the Arctic Ocean. Target sequences were generated as gBlocks (Integrated DNA Technologies, IA, USA) (Data S2), used in standard serial dilutions (10^0^-10^9^ gene copies), and tested against all primer-probe combinations to validate qPCR efficiency (Supporting Information Table S3) and specificity (i.e. confirm lack of non-target amplification). NCD assays were run in quadruplicate (DNA, 2.5 ng) or duplicate (cDNA, 1.5 μl; Beta-Arctic1, 3 μl) 10 μl reactions using a LightCycler® 480 Probes Master (Roche, Basel, Switzerland) with 1x LightCycler® 480 Probes Master mix, 0.5 μM primers, and 0.2 μM probe at 95°C-300 s, 40-45 cycles of 95°C-10 s, 60°C-30 s, and 72°C-1 s, and finally 40°C-10 s. UCYN-A and *Nodularia* sp. assays (UCYN-A1, UCYN-A2/A4, *Nodularia* sp.) (Supporting Information Table S3) were run in triplicate 12.5 μl reactions (Stratagene Mx3005) with 5 ng DNA or 0.8 ng cDNA, 1x Taqman® Universal PCR Master Mix (Applied Biosystems, MA, USA), 0.4 μM primers (UCYN-A1: 0.6 μM), 0.2 μM probe, and 0.1 μg μl^−1^ BSA at 50°C-120 s, 95°C-10 min, and 45 cycles of 95°C-30 s and 60°C-60 s. Duplicate ten-fold dilutions (10^1^-10^8^) of target cyanobacterial standards (GENEWIZ, Suzhou, China) were included in each run. No-template controls (PCR grade water) did not generate any amplification. Inhibition controls (duplicate for each environmental DNA and cDNA sample) with an added standard concentration of 10^6^ copies μl^−1^ showed no inhibition of amplification caused by template material (DNA or cDNA). Limits of detection (LOD) and quantification (LOQ) were established for each run (Armbruster and Pry 2008). Amplification below LOD was defined as 0 copies L^−1^, and amplification above LOD but below LOQ assigned a conservative number of 1 copy L^−1^. qPCR and RT-qPCR data are reported in Data S1.

### Statistical analysis

Statistical analyses were performed with R (v.4.1.0). R-packages ggplot2 (v.3.4.0; Wickham, 2016) and ggOceanMaps (v.1.3.4; Vihtakari, 2022) were used for data visualization. Initial pre-processing of *nifH* ASVs in phyloseq (v.1.36.0; McMurdie and Holmes, 2013) removed singletons (94 ASVs) for all downstream analyses except alpha diversity estimates based on singleton-containing median-normalized data. ASV count tables were center log-ratio transformed for principal component analysis (PCA; covariance matrix) using Aitchison distance (Gloor et al. 2017). The function envfit (vegan v.2.6-2; Oksanen et al. 2022) was used to explore relationships between continuous environmental variables and principal components.

Redundancy analysis (RDA) evaluated the proportion of variation explained by a selection of variables from a priori hypotheses and significant envfit output for DNA and cDNA, respectively. The final RDA model was evaluated on the top 50 ASVs, validated through the functions anova.cca and vif.cca (<20), and visualized with the R-package microViz (v.0.10.0; Barnett et al. 2021).

Categorical differences were evaluated by permutational multivariate analysis of variance with the function adonis2 on Aitchison distance matrices. To ensure homogeneity of group dispersions, the function permutest of betadisper output was used in parallel. Associations between ASVs and categorical variables were assessed with multipatt from the R-package indicspecies (v.1.7.12; Cáceres and Legendre, 2009). 999 permutations were used for permutational analyses (envfit, adonis2, permutest, betadisper, and multipatt).

Differences in *nifH* gene/transcript abundance (qPCR/RT-qPCR; log1p-transformed) were evaluated through Kruskal-Wallis with Dunn’s test for pairwise comparisons (with Bonferroni correction) between categorical variables. The relationship between continuous environmental variables (z-scored) and *nifH* gene/transcript abundance for each diazotroph group was explored through 1) Spearman’s rank correlation for monotonic relationships and visualized with corrplot (v.0.92; Wei and Simko, 2021), 2) generalized additive models for potential non-monotonic relationships, and 3) two-step generalized linear mixed model (GLMM) with a binary and a log-normal step to account for zero-heavy data (Thiele and Markussen 2012).

## Results

### Characterization of ecological water provinces

Sampling spanned 33 stations over ∼25 latitudinal degrees: in summer from the northern Norwegian Sea to the sea ice edge north of Svalbard, and in autumn from north of Svalbard to the northern Kattegat (mouth of the brackish Baltic Sea) (Fig. 1A). Water temperature ranged from -1.1 to 16.9°C and salinity ranged from 23.43 to 35.01 (Table 1; Supporting Information Fig. S3; Data S1). Based on the hydrographical and environmental conditions of the sub-surface water (using a continuously running underway water intake system), three ecological water provinces (hereafter denoted as “provinces”) were identified. A threshold salinity of 34.65 (Rudels et al. 2000) initially separating the northern fresher Arctic Water (ArW; north-western Barents Sea) from the saline Atlantic Water (AW; south-western Barents Sea) was applied and refined for small-scale variability in western Svalbard waters (Cottier et al. 2005). The same threshold was applied for the distinction between AW and fresher Coastal Influenced water to the south (CI; salinity <34.65; here referring to the Norwegian coastal current and the eastern North Sea/Skagerrak). The definition of the three provinces ArW, AW, and CI was consolidated through PCA of all environmental variables (Fig. 1B; Table 1), supporting the ecological relevance of the classification scheme used. Except for stations TR25-27 (TR denoting “transect” for all stations), which showed marginally (0.07-0.19) higher salinity than 34.65, their geographical location combined with the PCA results defined them as part of the CI province. ArW in the north (sampled both summer and autumn) was characterized by low inorganic nutrients, high fluorescence (summer), low temperature, and high HNA:LNA (a proxy for bacterial activity) (Fig. 1B; Table 1). AW (sampled both summer and autumn) in between ArW and CI was characterized by high salinity, high inorganic nutrient concentrations, and high molar N:P ratios ((NO_2_^−^+NO_3_^−^+NH_4_^+^)/PO_4_^3-^). CI in the south (sampled only in autumn) was characterized by low inorganic nutrient concentrations but higher bacterial, *Synechococcus,* and PPE abundances.

**Table 1.**
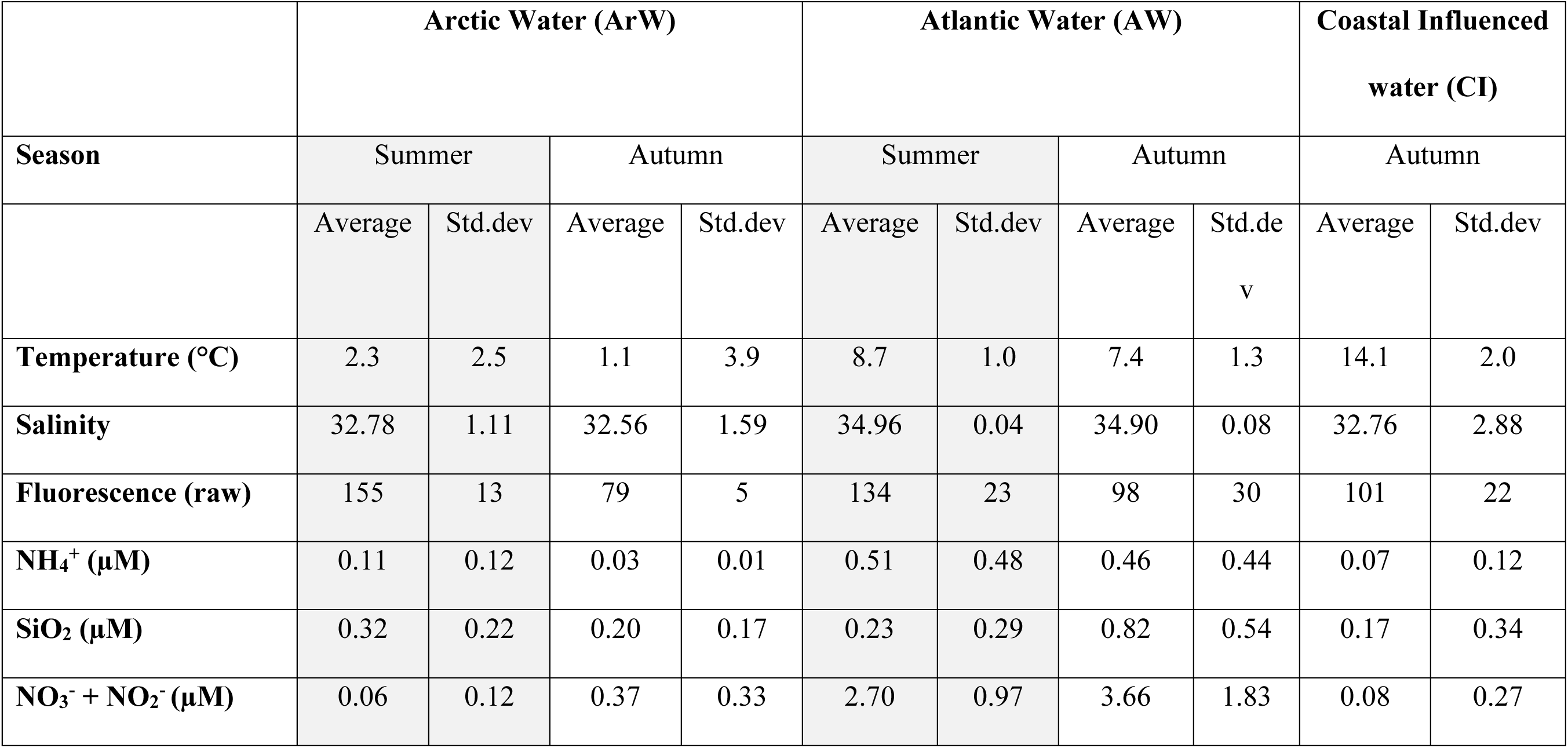
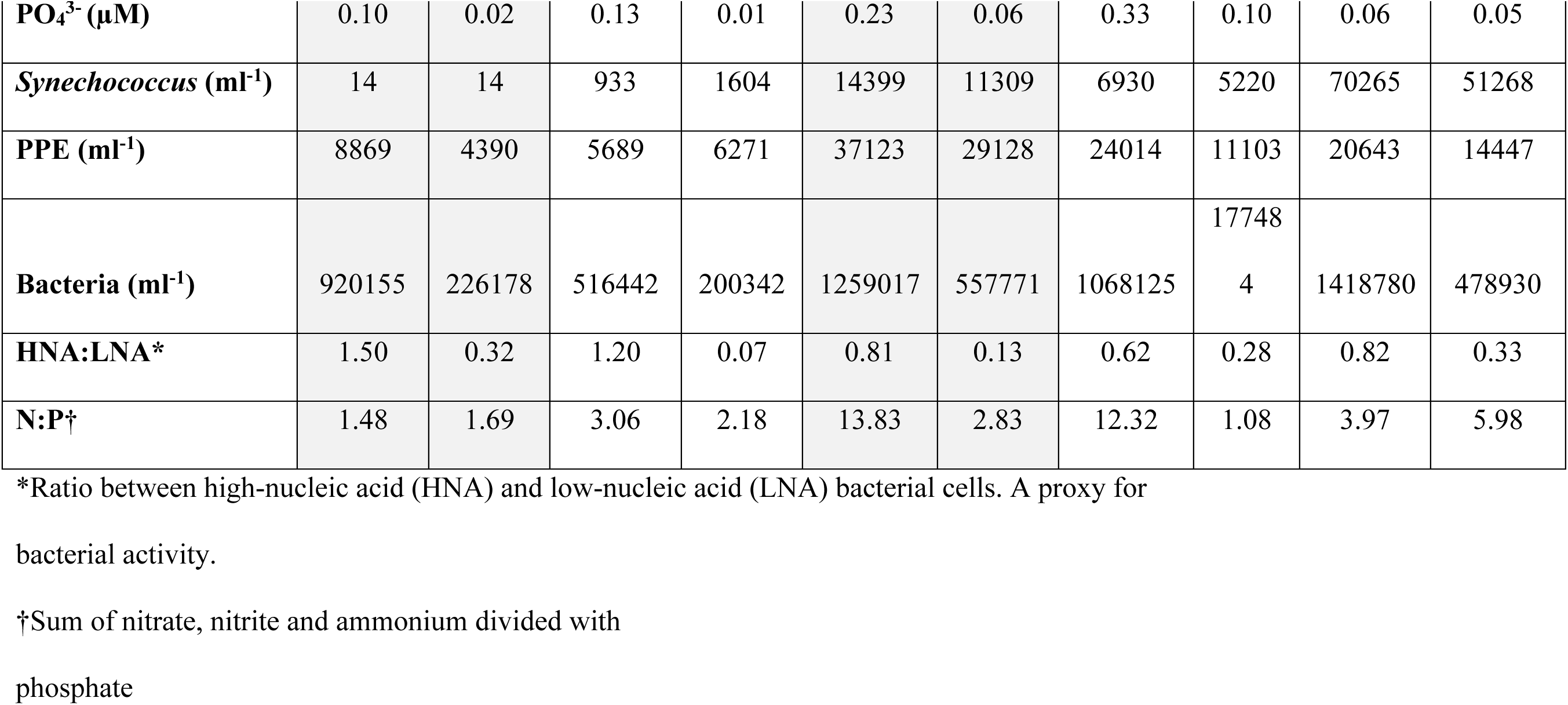
Environmental conditions in the different provinces. The average value and standard deviation (std.dev) for each variable in the different provinces are split into “summer” and “autumn”. Note that Coastal Influenced water (CI) was only sampled in autumn. PO_4_^3-^: phosphate, NO_3_^−^ + NO_2_^−^: nitrate + nitrite, NH_4_^+^: ammonium, SiO_2_: silicate, PPE: photosynthetic picoeukaryotes. Fluorescence is seen as a proxy for chlorophyll-*a*. HNA:LNA is the ratio between high-nucleic acid (HNA) and low-nucleic acid (LNA) bacterial cells and is a proxy for bacterial activity. N:P is the sum of nitrate, nitrite, and ammonium divided by phosphate (molar ratio).

|  | Arctic Water (ArW) |  |  |  | Atlantic Water (AW) |  |  |  | Coastal Influenced water (CI) |  |
| --- | --- | --- | --- | --- | --- | --- | --- | --- | --- | --- |
| Season | Summer |  | Autumn |  | Summer |  | Autumn |  | Autumn |  |
|  | Average | Std.dev | Average | Std.dev | Average | Std.dev | Average | Std.de<br>v | Average | Std.dev |
| Temperature (°C) | 2.3 | 2.5 | 1.1 | 3.9 | 8.7 | 1.0 | 7.4 | 1.3 | 14.1 | 2.0 |
| Salinity | 32.78 | 1.11 | 32.56 | 1.59 | 34.96 | 0.04 | 34.90 | 0.08 | 32.76 | 2.88 |
| Fluorescence (raw) | 155 | 13 | 79 | 5 | 134 | 23 | 98 | 30 | 101 | 22 |
| $\text{NH}_4^+$ (μM) | 0.11 | 0.12 | 0.03 | 0.01 | 0.51 | 0.48 | 0.46 | 0.44 | 0.07 | 0.12 |
| $\text{SiO}_2$ (μM) | 0.32 | 0.22 | 0.20 | 0.17 | 0.23 | 0.29 | 0.82 | 0.54 | 0.17 | 0.34 |
| $\text{NO}_3^- + \text{NO}_2^-$ (μM) | 0.06 | 0.12 | 0.37 | 0.33 | 2.70 | 0.97 | 3.66 | 1.83 | 0.08 | 0.27 |
| **PO<sub>4</sub><sup>3-</sup> (μM)** | 0.10 | 0.02 | 0.13 | 0.01 | 0.23 | 0.06 | 0.33 | 0.10 | 0.06 | 0.05 |
| ***Synechococcus* (ml<sup>-1</sup>)** | 14 | 14 | 933 | 1604 | 14399 | 11309 | 6930 | 5220 | 70265 | 51268 |
| **PPE (ml<sup>-1</sup>)** | 8869 | 4390 | 5689 | 6271 | 37123 | 29128 | 24014 | 11103 | 20643 | 14447 |
| **Bacteria (ml<sup>-1</sup>)** | 920155 | 226178 | 516442 | 200342 | 1259017 | 557771 | 1068125 | 17748<br>4 | 1418780 | 478930 |
| **HNA:LNA\*** | 1.50 | 0.32 | 1.20 | 0.07 | 0.81 | 0.13 | 0.62 | 0.28 | 0.82 | 0.33 |
| **N:P†** | 1.48 | 1.69 | 3.06 | 2.18 | 13.83 | 2.83 | 12.32 | 1.08 | 3.97 | 5.98 |

\*Ratio between high-nucleic acid (HNA) and low-nucleic acid (LNA) bacterial cells. A proxy for
bacterial activity.
†Sum of nitrate, nitrite and ammonium divided with
phosphate

Prokaryotic community composition (16S rRNA gene amplicon sequencing) also reflected the defined water provinces, supporting the ecological meaning of the categorization (Fig. 2A). Differences between provinces were, for example, evident as high relative abundances of Cyanobacteria in CI (39.7 ± 27.6%) and of Bacteroidota in ArW (27.7 ± 17.2%) (Fig. *2*B). Cyanobacterial reads were absent north of 80.352°N (station TR08 in the ArW) (Fig. *2*B) and flow cytometry counts revealed that *Synechococcus* was absent north of 79.426°N (station TR17 in the ArW) (Supporting Information Fig. S3).

### Diazotroph composition across environmental gradients

Diazotroph community composition (*nifH* amplicon sequencing) differed between the provinces (adonis2, *R^2^*=0.08, *p*=0.001, nested for season) (Fig. 3A-B). Cyanobacterial diazotrophs had high relative abundances in CI (29.3 ± 24.6%) but were sparse in AW and ArW (0.20 ± 0.37 and 0.08 ± 0.12%, respectively). In contrast to cyanobacterial diazotrophs, Delta- and Betaproteobacterial diazotrophs featured higher relative abundances in AW (21.0 ± 37.9% and 4.2 ± 17.9%, respectively) and ArW (20.0 ± 41.4% and 9.65 ± 29.8%, respectively), than in CI (0.83 ± 3.40 and 0.06 ± 0.11%, respectively) (Fig. 3A). Diazotroph alpha diversity (Shannon and Chao1) was lower in AW than in ArW and CI (Kruskal Wallis, df=2, *χ^2^*_shannon_=25.49, *χ^2^*_chao1_=12.42, *p*<0.01) (Supporting Information Fig. S4). *nifH* sequences are phylogenetically grouped into clusters (Chien and Zinder 1996), and here Cluster 1 dominated across all provinces, but a few stations in ArW and AW were dominated by Cluster 3, and stations TR12 and TR13 in ArW had a high contribution from Cluster 2 (>85%; vanadium nitrogenase, *vnf*) (Supporting Information Fig. S5). The variation in diazotroph community composition could be to 28.6% explained by temperature, salinity, fluorescence, HNA:LNA, NH_4_^+^, NO ^−^ and PO_4_^3-^ through Redundancy Analysis (RDA, *F*=4.23, *p*=0.001) (Fig. 3B; see Fig. 4 for the phylogenetic placement of ASVs).

**Fig. 4.**
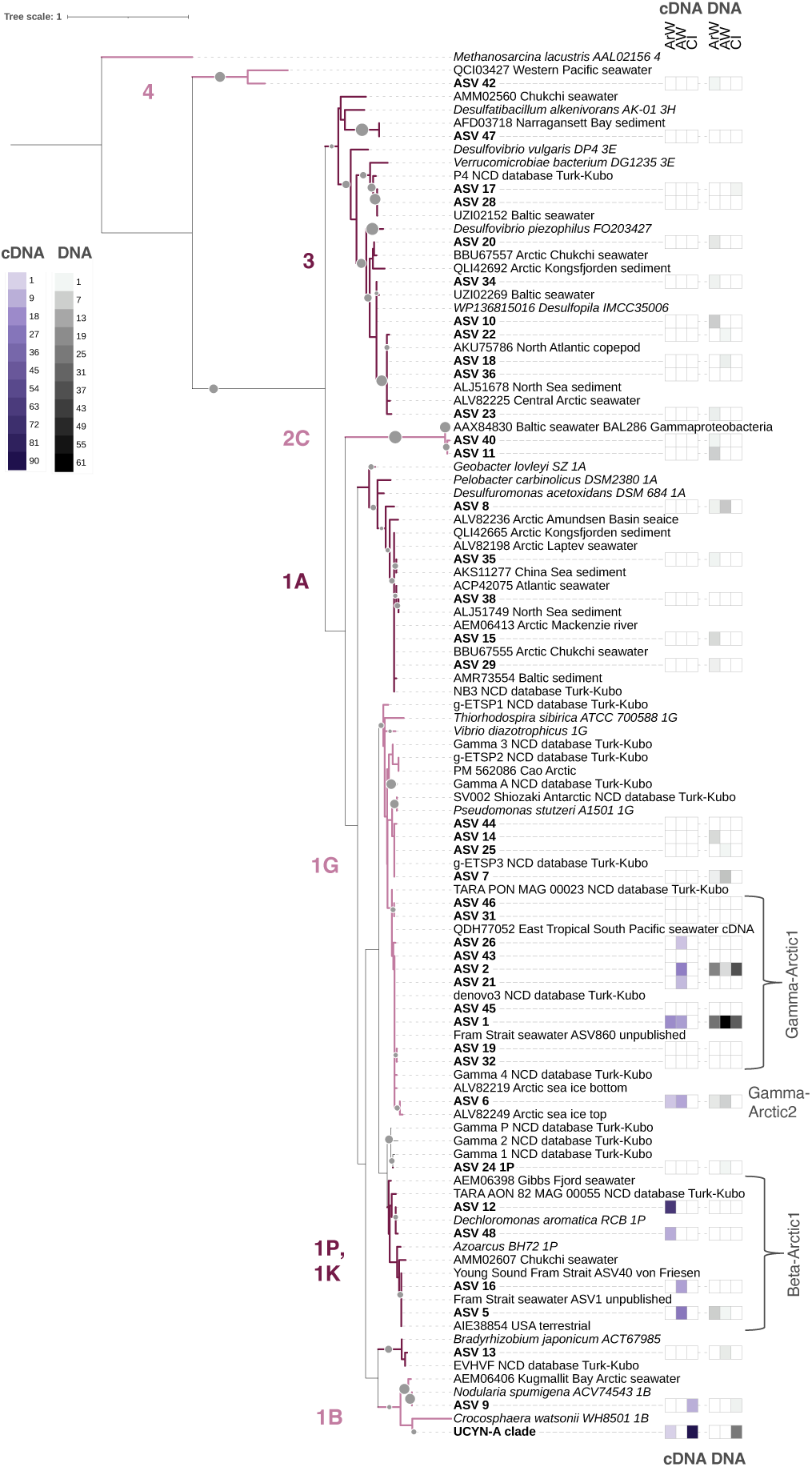
Partial *nifH* maximum-likelihood phylogenetic tree (amino acid sequences) of the top 50 amplicon sequence variants (ASVs) and reference sequences. *nifH* clusters/subclusters are denoted on the left side of the tree (Cluster 4: likely non-functional nitrogenases; Cluster 3: putative anaerobes, here mainly Desulfovibrionales and Desulfobacterales from Subcluster 3E; Subcluster 2C: alternative nitrogenase (*vnf*); 1A: here mainly Desulfuromonadales; 1G: Gammaproteobacteria; 1P and K: here mainly Rhodocyclales; 1B: Cyanobacteria). The accession number and origin of environmental reference sequences are denoted in each node label (e.g. “BBU67555 Arctic Chukchi seawater”). Sequences from the NCD catalog (Turk-Kubo et al. 2022) are marked with “NCD database Turk-Kubo” in its node label. Sequences from von Friesen et al. (2023) are indicated with “von Friesen”. Bootstrap values based on 100 iterations are presented as grey scaled circles from 50-100%. The heat maps display the relative abundance of each ASV in the provinces ArW (Arctic Water), AW (Atlantic Water), and CI (Coastal Influenced Water) for cDNA (left) and DNA (right), respectively. The three novel qPCR primer/probe sets targeting transcribing ASVs in AW and ArW (Supporting Information Table S3, Data S2) are denoted as “Gamma-Arctic1”, “Gamma-Arctic2” and “Beta-Arctic1” to the right of the tree. Note that all UCYN-A sublineages are collapsed into one clade. See Supporting Information Fig. S6A-B for the relative abundance of each UCYN-A sublineage and NCD group across the transects.

### Active diazotrophs across environmental gradients

From station TR22 (AW) southward to station TR23 (CI) in the south-western Barents Sea, *nifH*-transcription (i.e. active diazotrophic communities from *nifH* cDNA amplicon sequencing) changed from NCDs to UCYN-A (Fig. 3C). The transition was associated with a decline in inorganic nutrient concentrations and salinity (Supporting Information Fig. S3) and represents the province switch from AW to CI (Fig. 1A-B). The CI province thus contained different *nifH* transcripts compared to the northward provinces AW and ArW (adonis2, *R^2^*=0.17, *p*=0.001), represented by UCYN-A1, UCYN-A2, UCYN-A4, and *Nodularia* sp. (Supporting Information Fig. S6B). In contrast to CI, the actively *nifH*-transcribing diazotrophs in the ArW and AW were Gammaproteobacteria (Subcluster 1G) and Betaproteobacteria (Subcluster 1P) (Fig. 3C, Fig. **4**, Supporting Information Fig. S6B). The detected *nifH* transcripts along the autumn transect were all affiliated with Cluster 1 (Supporting Information Fig. S5B). Influence by temperature, PO_4_^3-^, and NO_3_^−^ was suggested, with a combined 21.8% variance explained for *nifH* transcript composition (RDA, *F*=1.58, *p*=0.001) (Fig. 3D; see Fig. 4 for the phylogenetic placement of ASVs). In summer, *nifH* transcripts could only be amplified from station TR02 in the AW (Fig. 3C, S4B) despite extensive PCR troubleshooting.

### Distribution and activity of non-cyanobacterial diazotrophs

North of the CI/AW transition in autumn, the active (i.e. *nifH* transcription) NCDs consisted of three phylogenetically distinct Beta- and Gammaproteobacterial groups (10 ASVs) (Fig. 4; Data S2). For these, qPCR assays were developed (“Beta-Arctic1”, “Gamma-Arctic1”, “Gamma-Arctic2”) (Supporting Information Table S3) to enumerate group-specific abundances.

The Gamma-Arctic1 ASVs (Subcluster 1G) were detected in all provinces (Supporting Information Fig. S6A) but almost exclusively transcribed in AW and ArW (Fig. 4, Fig**. 5**A, Supporting Information Fig. S6B). Gamma-Arctic1 is distinct from the Pacific Gamma4 (85.6% nucleotide similarity; Cheung et al. 2021), originally described in Halm et al. (2012). Gamma-Arctic1 is also distinct from the widely reported GammaA (79.8%, originally described in Moisander et al. (2008). However, Gamma-Arctic1 is similar (99.1-100%) to sequences previously reported from Arctic (Fernández-Méndez et al. 2016), North Pacific (“denovo3”; Gradoville et al. 2020), and mesopelagic subtropical waters (“9033127”; Salazar et al. 2019), and to transcripts in the eastern tropical South Pacific (Chang et al. 2019) (Fig. 4). Gamma-Arctic1 were quantifiable with qPCR throughout the autumn transect with a maximum of 8.2 × 10^3^ *nifH* gene copies L^−1^ in the AW province (station TR22) (Fig. 5A), which agrees with the *nifH* amplicon results (Supporting Information Fig. S6A). Gamma-Arctic1 gene copies correlated positively with temperature in a two-step generalized linear mixed model (GLMM, *p*=0.02) and negatively with fluorescence (GLMM, *p*=0.02) (Fig. 6, Supporting Information Fig. S7). Gamma-Arctic1 *nifH* transcripts never exceeded the limit of reverse RT-qPCR detection (85 copies L^−1^) (Supporting Information Table S3).

**Fig. 5.**
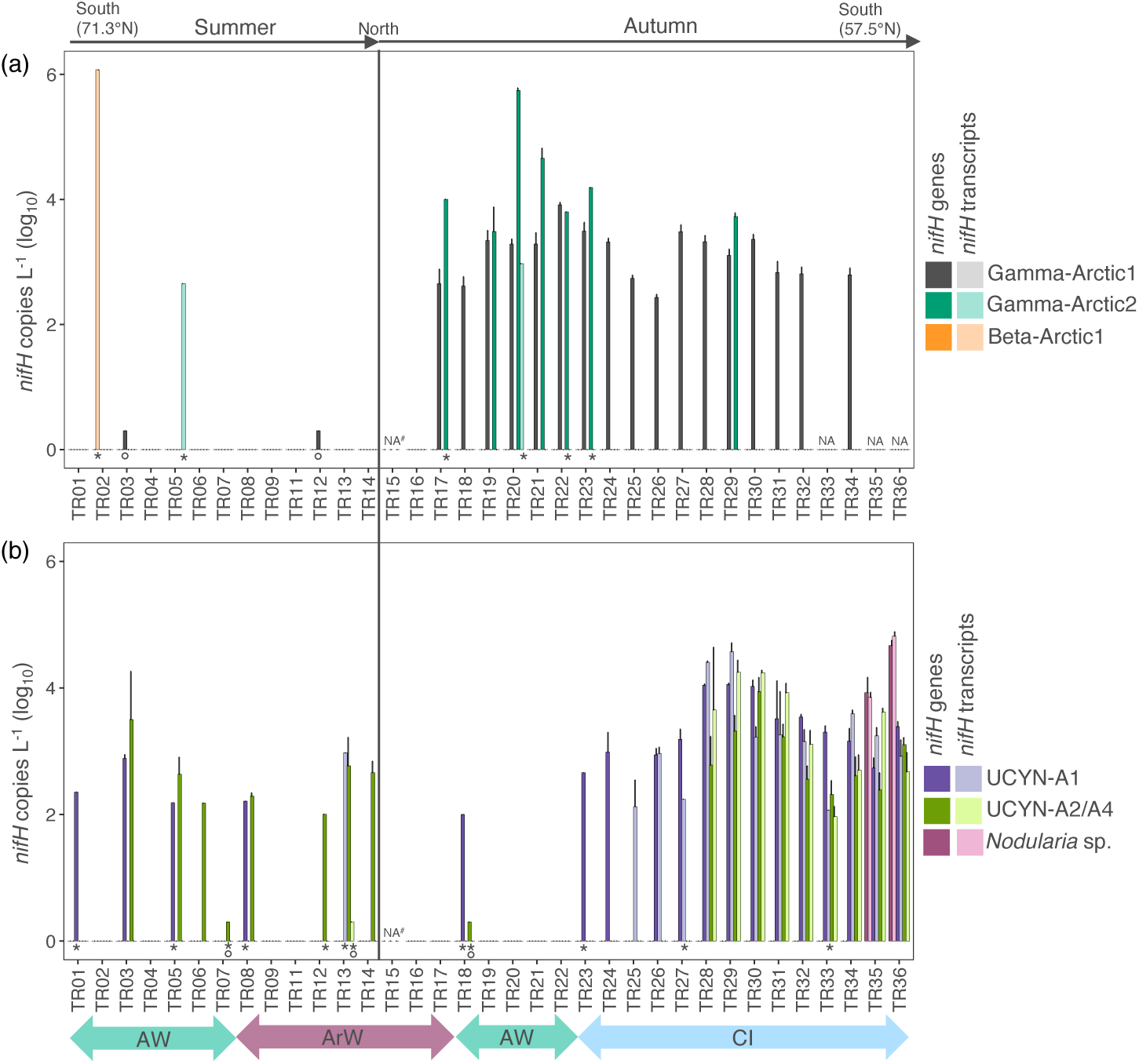
*nifH* gene and transcript copies along the summer and autumn transects assessed by quantitative PCR. **(a)** Non-cyanobacterial diazotrophs (Gamma-Arctic1, Gamma-Arctic2 and Beta-Arctic1) and **(b)** UCYN-A1, UCYN-A2/A4, and *Nodularia* sp. Error bars represent standard deviation. Limit of detection (LOD) and quantification (LOQ), qPCR assay efficiency (%), R^2^ (standard curve correlation coefficient of determination), and targeted amplicon sequence variants are specified in Supporting Information Table S3. ArW: Arctic Water, AW: Atlantic Water, CI: Coastal Influenced Water. *: only one replicate amplified, °: above LOD but below LOQ. NA: not analyzed. NA^#^: only DNA analyzed. No bar: analyzed but not detected.

**Fig. 6.**
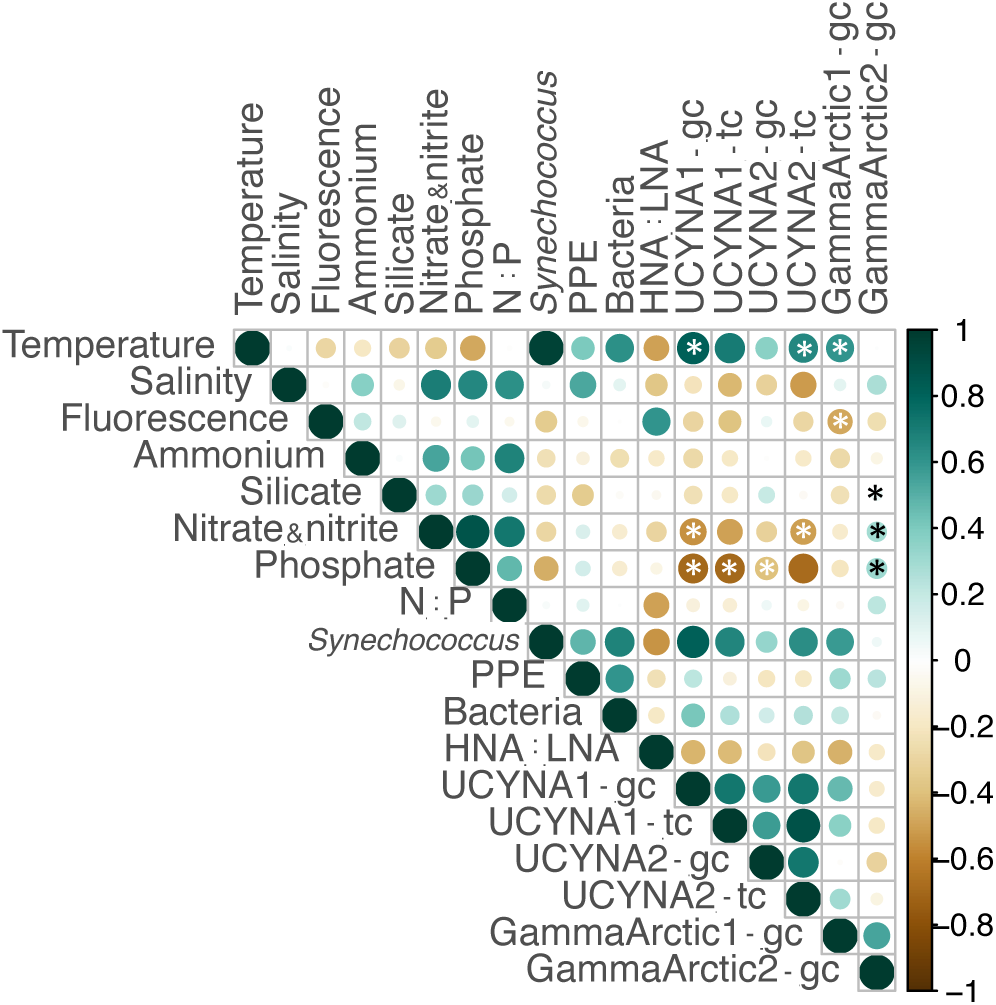
Correlations of *nifH*-gene and transcripts with environmental variables. Spearman rank correlation matrix of *nifH* gene- and transcript copies L^−1^ with environmental variables for the diazotrophs targeted with quantitative PCR (>2 observations). N:P: inorganic nitrogen to phosphorus molar ratio, PPE: photosynthetic picoeukaryotes, HNA:LNA: high nucleic acid to low nucleic acid bacterial cell ratio (a proxy for bacterial activity), gc: gene copies, tc: transcript copies. Asterisks denote significant correlation from two-step generalized linear mixed models (GLMM).

Gamma-Arctic2 (Subcluster 1G; ASV6) was associated with ArW and AW (multipatt, *p*=0.008) (Fig. 3B, Fig**. 3**D, Fig**. 4**) and is 97.6% similar to a diazotroph reported from Arctic sea ice (Fernández-Méndez et al. 2016), and 87.0% similar to the Pacific Gamma4 (Cheung et al. 2021) (Fig. Fig**. 4**). Further, it is 95.4% similar to *Amphritea atlantica* (AP025284) – an aerobic, motile, organic matter degrading Gammaproteobacterium isolated from hydrothermal vent bivalves (Gärtner et al. 2008). Gamma-Arctic2 was mainly found in AW in autumn (up to 5.5 × 10^5^ *nifH* gene copies L^−1^) (Fig. 5A), and was positively correlated with inorganic nutrients (GLMM, *p*_PO4_=0.0030; p_NO3_=0.0004; *p*_SiO2_=0.0021) (Fig. 6, Supporting Information Fig. S7). Gamma-Arctic2 *nifH* transcripts were quantifiable with RT-qPCR exclusively at nitrogen-replete (>4 μM NO_3_^−^+NO_2_^−^) locations (TR05: 4.5 × 10^2^ copies L^−1^; TR20: 9.4 × 10^2^ copies L^−1^) (Fig. 5A).

Beta-Arctic1 (Subcluster 1P) ASVs were primarily detected in the ArW and AW (Fig. 4 and Supporting Information Fig. S6) and are affiliated with the order Rhodocyclales, previously reported from the Arctic regions of the Chukchi Sea (Shiozaki et al. 2018), and Fram Strait and Young Sound (von Friesen et al. 2023). Beta-Arctic1 had 98.2-100% amino acid similarity to *Dechloromonas aromatica* (Q47G67.1) (Fig. 4) – a facultative anaerobic Betaproteobacterium known from soil and freshwater, capable of biofilm formation, chemotaxis, degradation of aromatic compounds, and possibly associated to a eukaryotic host (Salinero et al. 2009). *nifH* gene copies of Beta-Arctic1 were below the limit of qPCR detection at all locations (137 *nifH* copies L^−1^), but transcripts were detected with RT-qPCR in AW in summer (station TR02; 1.2 × 10^6^ *nifH* copies L^−1^) (Fig. 5A). This number should be cautiously interpreted as only half of the replicates amplified, but Beta-Arctic1 was nevertheless confirmed in the *nifH* amplicon cDNA library from a separate replicate sample at this station (ASV 5; accounting for >95% relative abundance) (Supporting Information Fig. S6B), thus supporting Beta-Arctic1 activity.

### Distribution and activity of UCYN-A and *Nodularia* sp

Cyanobacterial diazotrophs were almost exclusively accounted for by UCYN-A (93%) (Fig. 4, Supporting Information Fig. S6). Among the top 100 ASVs, 20 ASVs were assigned as UCYN-A1, three as UCYN-A2, and one as UCYN-A4 (Supporting Information Fig. S2). UCYN-A and *Nodularia* sp. were also detected in the 16S rRNA gene libraries, with UCYN-A accounting for 0.01-0.09% relative abundance at stations TR28–32 in the CI, and *Nodularia* sp. for 0.03% and 0.15% at stations TR35 and TR36 in the CI, respectively.

Abundance and transcription patterns of *nifH* from *Nodularia* sp., UCYN-A1, and UCYN-A2/A4 were quantified with qPCR and RT-qPCR. Note that the applied primer/probe set cannot distinguish between sublineages UCYN-A2 and UCYN-A4 (Farnelid et al. 2016). In CI, *nifH* gene copies L^−1^ were up to 10^4^ for UCYN-A1 and *Nodularia* sp., and 10^3^ for UCYN-A2/A4 (Fig. 5B). UCYN-A1 and UCYN-A2/A4 *nifH* gene abundances were patchy in ArW and AW, whereas in CI, genes and transcripts (10^2^-10^4^ *nifH* copies L^−1^) frequently co-occurred. The UCYN-A sublineages overlapped southwards from TR28 in the CI, whereas only transcripts of UCYN-A1 were detected north of this station, as confirmed in both the RT-qPCR and cDNA *nifH* amplicon datasets (Fig. 5B, Supporting Information Fig. S6B). Transcripts from both UCYN-A1 and UCYN-A2/A4 were also detected at station TR13 in the marginal ice zone (-0.3°C; ArW province) north of Svalbard, where UCYN-A1 was also reported in the *nifH* gene amplicon library (Supporting Information Fig. S6A). UCYN-A4 transcription was detected (cDNA *nifH* amplicon sequencing) at stations TR28 and TR29 in the CI in the southern Norwegian Sea (Supporting Information Fig. S6B).

UCYN-A1 gene copies were positively correlated with temperature (GLMM, *p*=0.002) and negatively correlated with NO_3_^−^ (GLMM, *p*=0.003) and PO_4_^3-^ (GLMM, *p*=0.0005) (Fig. 6, Supporting Information Fig. S7). Analogously, UCYN-A1 transcript copies were negatively correlated with PO_4_^3-^ (GLMM, *p*=0.003). UCYN-A2/A4 gene copies were similarly negatively correlated with PO_4_^3-^ (GLMM, *p*=0.006), whereas transcripts were only quantifiable where NO_3_^−^ concentrations were <0.01 μM (GLMM, *p*=0.001) and positively correlated with temperature (GLMM, *p*=0.0131) (Fig. 6, Supporting Information Fig. S7). UCYN-A sublineages displayed maximum transcript abundances at ∼14°C (stations TR28–30 in the CI) (Fig. 5B, Supporting Information Fig. S7). UCYN-A *nifH* genes and transcripts were positively correlated with *Synechococcus* abundance (Fig. 6). Presence and transcription of *Nodularia* sp. *nifH* genes were observed at the two stations closest to the mouth of the Baltic Sea (TR35 and TR36; salinity 32.99 and 23.43; 10^3^ and 10^4^ copies L^−1^, respectively) (Fig. 5B), consistent with the *nifH* amplicon library findings (Supporting Information Fig. S6A-B).

## Discussion

In this study, we show that diazotrophs with contrasting metabolisms (i.e. photoautotrophs or chemoheterotrophs) differ markedly in biogeography between temperate-boreal and polar waters. This was established based on the sharp transition from photosynthesis-based diazotrophy (UCYN-A) in the North Atlantic to probable chemoheterotrophic (non-cyanobacterial) diazotrophy in the gateway of the Arctic Ocean. Upon entering nutrient-rich waters in the Atlantic gateway to the Arctic Ocean, the abrupt transition in *nifH* transcription indicates inorganic nutrients as probable drivers of this distinction between polar and temperate-boreal diazotroph communities and their activity.

### Non-cyanobacterial diazotrophy dominates the Atlantic gateway

We propose a prominent role of NCDs in polar waters. Resembling indications in the Pacific gateway to the Arctic Ocean, we here report a transition to non-cyanobacterial diazotrophy in the Atlantic gateway to the Arctic. The observed dominance of NCDs upon entering Atlantic-derived water in the Barents Sea and the three active (*nifH* transcribing) NCD groups (Beta-Arctic1: Rhodocyclales, Gamma-Arctic1: unknown, Gamma-Arctic2: Oceanospirillales) suggest a prominent role of NCDs in these Arctic waters. Identical or similar phylotypes to the herein-identified key NCDs Beta-Arctic1, Gamma-Arctic1, and Gamma-Arctic2, have previously been reported to occur in different regions of the Arctic Ocean: the coastal Canadian Arctic (Blais et al. 2012), Central Arctic sea ice (Fernández-Méndez et al. 2016), Chukchi Sea (Shiozaki et al. 2018), and Fram Strait and Young Sound (von Friesen et al. 2023). Now backed up by *nifH* transcription data in the present study (although it should be noted that not all RT-qPCR replicates amplified), we propose that they are key diazotrophs in the Atlantic-influenced Arctic Ocean and possibly also more broadly across the Arctic. Their quantitative contribution to nitrogen fixation remains to be elucidated, but the functional composition of polar diazotrophs is distinct from temperate-boreal waters.

Biogeographical distributions of NCDs are phylotype-specific. The three Arctic NCD groups exhibited different biogeographical distributions, with Beta-Arctic1 and Gamma-Arctic2 being associated with the polar region (AW and ArW), whereas genes (but not transcripts) representing Gamma-Arctic1 were also detected in the temperate-boreal region (CI). Combined with findings of Gamma-Arctic1 in e.g. subtropical mesopelagic waters (Salazar et al. 2019), our data suggest that Gamma-Arctic1 has a more widespread distribution. In contrast, the closest related environmental sequences to Gamma-Arctic2 and Beta-Arctic1 were from other Arctic regions (Blais et al. 2012; Fernández-Méndez et al. 2016; von Friesen et al. 2023), suggesting that they may mostly or exclusively be associated with the polar region. The pattern with NCD phylotypes exhibiting either an Arctic-associated or more widespread distribution is consistent with patterns that have been identified in Arctic metagenome-assembled genomes, although with low sequence similarity to the here-discussed NCDs (<77%; Shiozaki et al. 2023). Endemism has recently been suggested for several Arctic NCDs from the Canadian Arctic gateway (outflow region of the Arctic Ocean; Robicheau, Tolman, Rose, et al. 2023). Taken together, some NCDs active in the Arctic are specific to the polar biome, whereas others are widespread across multiple ocean regions.

### UCYN-A in the northern North Atlantic Current

We document a yet wider distribution of UCYN-A sublineages in the global ocean. We frequently observed and quantified active UCYN-A sublineages primarily in the CI province (along the Norwegian coast) with a more irregular presence further north in the polar region (AW and ArW). Interestingly, transcription of both UCYN-A1 and UCYN-A2 were detected in the marginal ice zone (in the ArW) with identical *nifH* ASVs to those further south in the CI, similar to findings from the Pacific Arctic (Harding et al. 2018). It should be noted, however, that a recently constructed Arctic UCYN-A2 metagenome-assembled genome held specific adaptations despite having a *nifH* sequence identical to that of UCYN-A2 from non-polar regions (Shiozaki et al. 2023). Hence, it cannot be excluded but rather expected that identical UCYN-A2 phylotypes found along the transects can differ in adaptive traits and ecological features.

We speculate that the marginal ice zone may be an advantageous habitat for UCYN-A and its host due to sea ice melt containing elevated iron concentrations (Aguilar-Islas et al. 2008), which is required for both nitrogenase and photosynthesis systems. In support of this, UCYN-A *nifH* gene abundance was recently reported to correlate with a high ratio of iron to inorganic nitrogen (Fe:N) in the vicinity of a river in the low Pacific Arctic (Cheung et al. 2022). The findings of active UCYN-A in the marginal ice zone in our study may, due to its relatively coastal location at the time of sampling, be boosted by a combination of sea ice and river-derived iron. Measurements of iron concentration and speciation are thus encouraged in future studies.

An observed correlation between UCYN-A and *Synechococcus* could indicate ecological interaction, as the UCYN-A host is thought to prey on *Synechococcus* (Frias-Lopez et al. 2009) or may indicate similar environmental niches among them. Having previously been reported to co-occur in the coastal north-west Atlantic (Robicheau et al. 2023a), this merits further study. Overall, we report UCYN-A from the Norwegian and Barents Seas – expanding their known biogeographical range (Farnelid et al. 2016), which will enable refined distribution modeling.

### Nutrients as potential key drivers of diazotrophy

Inorganic nitrogen availability appears to be an important regulator for the relative abundance of different diazotroph groups. Transcription of *nifH* by UCYN-A2/A4 could, contrary to recent findings (Selden et al. 2022), only be quantified at nitrogen-depleted locations, implying regulation by inorganic nitrogen or potential co-varying parameters. Interestingly, the transition to non-cyanobacterial diazotrophy corresponded with a marked increase in inorganic nitrogen concentration (especially for Gamma-Arctic2 and Beta-Arctic1). This is in agreement with reports of NCD activity from other nitrogen-replete waters (Knapp 2012) and in culture (Bentzon-Tilia et al. 2015), suggesting an inverse relationship with inorganic nitrogen. The ability to bring inorganic and organic nitrogen compounds into the cell varies among NCDs (Turk-Kubo et al. 2022) and may explain the activity of NCDs in nitrogen-replete conditions in Arctic waters. Related to this, a switch from nitrogen fixation to nitrate uptake and reduction can even be energetically unfavorable under certain conditions (Karl et al. 2002). It is also proposed that nitrogen fixation may take place to ensure an intracellular redox balance rather than satisfying cellular nitrogen requirements (Bombar et al. 2016). The NO_3_^−^ concentration decreased again northwards from AW to ArW but was still 4.6 times higher (μM) in ArW compared to CI (autumn). This may have constrained potential UCYN-A activity in the ArW. There are likely several other important drivers not measured in the current study, such as grazing pressure (Deng et al. 2023) and micronutrient availability (e.g. iron; Knapp et al. 2016). While the underlying causal ecophysiological mechanisms remain to be explained, inorganic nitrogen is likely playing a role in governing diazotroph community composition and activity and, thus, the potential for nitrogen fixation.

Phytoplankton aggregates, including those originating from diatoms, may provide accessible carbon and foster low oxygen conditions that are favorable for NCD nitrogen fixation (Riemann et al. 2022). In our study, the *nifH* gene abundance of Gamma-Arctic1 was negatively correlated with fluorescence. This contrasts with the Gammaproteobacterial NCD Gamma4 in the North Pacific being positively correlated with chlorophyll-*a* (Cheung et al. 2021) and suggests different associations between Gammaproteobacterial NCDs and phytoplankton biomass. The positive correlation of Gamma-Arctic2 with dissolved silicate resembles patterns of the widespread GammaA in tropical waters where a positive correlation to silicate was proposed as a potential association to diatoms (Shao and Luo 2022), further consistent with its suggested particle-associated lifestyle (Cornejo-Castillo and Zehr 2021). GammaA was recently confirmed as an actively nitrogen-fixing diatom symbiont of alphaproteobacterial origin that has likely obtained the nitrogenase enzyme complex through horizontal gene transfer from a Gammaproteobacterium (Tschitschko et al. 2024). This finding expands known diazotroph-diatom associations to also include NCDs and encourages future metagenomic analyses to supplement *nifH*-inferred phylogeny. Conclusively, it remains elusive whether the correlations between the here reported Gamma-Arctic2 and silicate may indicate an indirect (through particulate and/or dissolved organic matter provision) or a direct (through potential facultative or obligate symbiosis) link to diatoms.

Our findings imply a markedly broader salinity tolerance of the filamentous, heterocystous Cyanobacteria *Nodularia* sp. than previously assumed. *Nodularia* sp. is a key diazotroph in the Baltic Sea with a biomass peak around salinity 10 (Lehtimaki et al. 1997). Here, we report their *nifH* transcription at 2-3-fold higher salinities in the CI (32.99 and 23.43), suggesting a broader range.

### Seasonality

The influence of seasonality on northern (i.e. temperate-boreal and polar) diazotroph community composition and *nifH* transcription is poorly understood, with most studies being conducted in summer (von Friesen and Riemann 2020). It is unknown whether the switch from UCYN-A to NCDs at the CI to AW transition also occurred in summer as the southernmost sampling point was within the AW (i.e. CI was not sampled in summer in this study). The only previously reported annual data is within the Arctic region, preventing direct comparison (von Friesen et al. 2023). Within AW and ArW (sampled both summer and autumn), Gamma-Arctic1 and Gamma-Arctic2 occurred mainly in autumn, resembling seasonal patterns reported for Gamma4 in the North Pacific (Cheung et al. 2021). In contrast, UCYN-A was more abundant in summer similar to observations in other regions (Cabello et al. 2020). As UCYN-A is previously reported from several Arctic seas (Shiozaki et al. 2017; Harding et al. 2018), including outflow regions and relatively confined coastal areas (e.g. Robicheau, Tolman, Rose, et al. 2023; von Friesen et al. 2023), it is likely, but not proven, that they can endure long periods of darkness (i.e. having over-wintering strategies). However, if UCYN-A indeed is mainly advected into the Arctic from adjacent oceans, lack of light may ultimately limit the potential of UCYN-A to spread further northwards through atlantification. Seasonal differences in the variable diazotroph communities, and ultimately the associated nitrogen fixation, are of high future research priority.

### Nitrogen fixation: from temperate-boreal to polar waters

We hypothesize that nitrogen fixation by NCDs and UCYN-A constitute local sources of nitrogen in the polar (ArW and AW) and temperate-boreal (CI) regions, respectively, and encourage their future quantification. Our findings of *nifH* transcription by NCDs in Atlantic-originating and Arctic waters in the Barents Sea (up to 9.4 × 10^2^ transcripts L^−1^; Gamma-Arctic2) suggest that they are actively fixing nitrogen. Previous NCD *nifH* gene abundances of similar magnitude as we report (10^2^-10^5^ copies L^−1^) have been paralleled with nitrogen fixation rates of 0.5-5.1 nmol N L^−1^ d^−1^ in the Eastern Tropical South Pacific in the absence of cyanobacterial diazotrophs (Gradoville et al. 2017). In comparison, UCYN-A *nifH* gene abundances of similar magnitude as we report along the transect (10^3^-10^4^ copies L^−1^) have been associated with nitrogen fixation rates of 0.004-3.6 nmol N L^−1^ d^−1^ in the Pacific Arctic (Shiozaki et al. 2018; Harding et al. 2018) and up to ∼6 nmol N L^−1^ d^−1^ in the North Pacific Subtropical Gyre (Gradoville et al. 2020). UCYN-A1 *nifH* transcript abundance in the marginal ice zone of our study was two orders of magnitude higher than in the Bering Sea (Pacific Arctic), where Shiozaki et al. (2017) measured nitrogen fixation rates of 2.8 nmol N L^−1^ d^−1^ in parallel to the *nifH* transcripts. While confirmation is needed through the direct measurement of nitrogen fixation, our results imply a high potential for UCYN-A and NCDs to contribute with new nitrogen to the Atlantic-influenced Arctic Ocean.

### Potential impacts of atlantification on diazotrophy

We speculate that the ecological boundary identified between *nifH*-expressing UCYN-A and NCDs in the south-western Barents Sea will shift northwards with continued atlantification. Here, we show that the transition from the warmer and nutrient-poor temperate-boreal Norwegian Sea to the colder and nutrient-rich Arctic Barents Sea has a major impact on diazotrophy. The changing hydrographical conditions brought on by atlantification impact organismal biogeography and are predicted to alter bacterial communities of the Arctic Ocean (Priest et al. 2023). These shifts may affect future nitrogen fixation potential and thereby influence associated primary productivity. It is likely that the here proposed key Arctic NCD groups, with either current Arctic-associated or more widespread distributions, will differ in their responses to a shrinking polar biome and that our findings are describing an already ongoing functional shift in diazotrophy. Considering the increasing primary production and nitrogen utilization, especially in the Atlantic-influenced Arctic Ocean (Lewis et al. 2020), the associated elevated inputs of labile organic matter from phytoplankton may stimulate NCDs. An increased advection of UCYN-A northward along the Norwegian Coastal Current, over the Barents Sea, and through the Fram Strait is a possible future scenario where their nitrogen fixation activity may be in part regulated by regional nutrient availability. With increased atlantification and expected elevated primary production and sea ice melt that can deliver additional iron and organic matter, we speculate that future conditions will become more favorable for diazotrophy. Quantification of nitrogen fixation on both sides of the herein-identified ecological boundary will be important to further constrain nitrogen budgets and refine predictions of primary production trajectories in the rapidly changing Arctic Ocean.

## Acknowledgments

Permission for sampling within Norwegian economic waters was given by the Norwegian Directorate of Fisheries (18.08.2021, 25.08.2021-23.09.2021, Jnr. 21/12086) and the Norwegian Petroleum Directorate (802/2021). The Swedish Polar Research Secretariat (SPRS; https://polar.se) organized and supported the SAS-Oden 2021 expedition with IB Oden in the Central Arctic Ocean. This expedition was the Swedish contribution to the International “Synoptic Arctic Survey” (SAS; https://synopticarcticsurvey.w.uib.no/). We thank the Master and crew of IB Oden for expertly undertaking the SAS-Oden 2021 expedition. We are grateful for laboratory assistance from Cecilie Appeldorff (University of Copenhagen), Andras Turai and Laura Bas Conn (Linnaeus University), and statistical guidance from Bo Markussen (University of Copenhagen). We thank Anna Willstrand Wranne (Swedish Meteorological and Hydrological Institute) for the use of equipment. The authors declare that they have no conflict of interest.

## Data availability

All data needed to evaluate the conclusions in the paper are present in the paper, the Supporting Information, the files Data S1 and Data S2 deposited at Zenodo (doi: 10.5281/zenodo.14652839), and the raw fastq files from *nifH* and 16S rRNA gene amplicon sequencing are deposited in the NCBI Sequence Read Archive under BioProject number PRJNA994547 and PRJNA994909, respectively (sample IDs listed in Data S1). Ship data from the Synoptic Arctic Survey expedition is available in the Swedish National Data Service [https://snd.gu.se/en/catalogue/study/2021-123]. Datasets from qPCR, RT-qPCR, sample IDs, and environmental conditions are available in Data S1. Non-cyanobacterial diazotroph *nifH* amplicon sequences for which qPCR primer/probe sets were designed are available in Data S2.

## References

Aguilar-Islas, A. M., R. D. Rember, C. W. Mordy, and J. Wu. 2008. Sea ice-derived dissolved iron and its potential influence on the spring algal bloom in the Bering Sea. Geophys. Res. Lett. 35: L24601. doi:10.1029/2008GL035736

Alegria Zufia, J., H. Farnelid, and C. Legrand. 2021. Seasonality of Coastal Picophytoplankton Growth, Nutrient Limitation, and Biomass Contribution. Front. Microbiol. 12: 1–13. doi:10.3389/fmicb.2021.786590

Angel, R., M. Nepel, C. Panhölzl, H. Schmidt, C. W. Herbold, S. A. Eichorst, and D. Woebken. 2018. Evaluation of Primers Targeting the Diazotroph Functional Gene and Development of NifMAP – A Bioinformatics Pipeline for Analyzing *nifH* Amplicon Data. Front. Microbiol. 9: 1–15. doi:10.3389/fmicb.2018.00703

Armbruster, D. A., and T. Pry. 2008. Limit of blank, limit of detection and limit of quantitation. Clin. Biochem. Rev. 29 Suppl 1: S49–52.

Årthun, M., T. Eldevik, L. H. Smedsrud, Ø. Skagseth, and R. B. Ingvaldsen. 2012. Quantifying the Influence of Atlantic Heat on Barents Sea Ice Variability and Retreat. J. Clim. 25: 4736–4743. doi:10.1175/JCLI-D-11-00466.1

Barnett, D., I. Arts, and J. Penders. 2021. microViz: an R package for microbiome data visualization and statistics. J. Open Source Softw. 6: 3201. doi:10.21105/joss.03201

Bentzon-Tilia, M., I. Severin, L. H. Hansen, and L. Riemann. 2015. Genomics and Ecophysiology of Heterotrophic Nitrogen-Fixing Bacteria Isolated from Estuarine Surface Water S.J. Giovannoni [ed.]. MBio 6: 1–11. doi:10.1128/mBio.00929-15

Blais, M., J. Tremblay, A. D. Jungblut, J. Gagnon, J. Martin, M. Thaler, and C. Lovejoy. 2012. Nitrogen fixation and identification of potential diazotrophs in the Canadian Arctic. Global Biogeochem. Cycles 26: GB3022. doi:10.1029/2011GB004096

Bombar, D., R. W. Paerl, and L. Riemann. 2016. Marine Non-Cyanobacterial Diazotrophs: Moving beyond Molecular Detection. Trends Microbiol. 24: 916–927. doi:10.1016/j.tim.2016.07.002

Boström, K. H., L. Riemann, U. L. Zweifel, and Å. Hagström. 2007. *Nodularia* sp. *nifH* gene transcripts in the Baltic Sea proper. J. Plankton Res. 29: 391–399. doi:10.1093/plankt/fbm019

Cabello, A. M., K. A. Turk-Kubo, K. Hayashi, L. Jacobs, R. M. Kudela, and J. P. Zehr. 2020. Unexpected presence of the nitrogen-fixing symbiotic cyanobacterium UCYN-A in Monterey Bay, California. J. Phycol. 56: 1521–1533. doi:10.1111/jpy.13045

Cáceres, M. De, and P. Legendre. 2009. Associations between species and groups of sites: indices and statistical inference. Ecology 90: 3566–3574. 10.1890/08-1823.1

Callahan, B. J., P. J. McMurdie, M. J. Rosen, A. W. Han, A. J. A. Johnson, and S. P. Holmes. 2016. DADA2: High-resolution sample inference from Illumina amplicon data. Nat. Methods 13: 581–583. doi:10.1038/nmeth.3869

Chang, B. X., A. Jayakumar, B. Widner, P. Bernhardt, C. W. Mordy, M. R. Mulholland, and B. B. Ward. 2019. Low rates of dinitrogen fixation in the eastern tropical South Pacific. Limnol. Oceanogr. 64: 1913–1923. doi:10.1002/lno.11159

Cheung, S., K. Liu, K. A. Turk-Kubo, and others. 2022. High biomass turnover rates of endosymbiotic nitrogen-fixing cyanobacteria in the western Bering Sea. Limnol. Oceanogr. Lett. 7: 501–509. doi:10.1002/lol2.10267

Cheung, S., J. P. Zehr, X. Xia, and others. 2021. Gamma4: a genetically versatile Gammaproteobacterial *nifH* phylotype that is widely distributed in the North Pacific Ocean. Environ. Microbiol. 23: 4246–4259. doi:10.1111/1462-2920.15604

Chien, Y. T., and S. H. Zinder. 1996. Cloning, functional organization, transcript studies, and phylogenetic analysis of the complete nitrogenase structural genes (*nifHDK2*) and associated genes in the archaeon *Methanosarcina barkeri* 227. J. Bacteriol. 178: 143–148. doi:10.1128/JB.178.1.143-148.1996

Church, M., B. Jenkins, D. Karl, and J. Zehr. 2005. Vertical distributions of nitrogen-fixing phylotypes at Stn Aloha in the oligotrophic North Pacific Ocean. Aquat. Microb. Ecol. 38: 3–14. doi:10.3354/ame038003

Coale, T. H., V. Loconte, K. A. Turk-Kubo, and others. 2024. Nitrogen-fixing organelle in a marine alga. Science (80-.). 384: 217–222. doi:10.1126/science.adk1075

Cornejo-Castillo, F. M., and J. P. Zehr. 2021. Intriguing size distribution of the uncultured and globally widespread marine non-cyanobacterial diazotroph Gamma-A. ISME J. 15: 124–128. doi:10.1038/s41396-020-00765-1

Cottier, F., V. Tverberg, M. Inall, H. Svendsen, F. Nilsen, and C. Griffiths. 2005. Water mass modification in an Arctic fjord through cross-shelf exchange: The seasonal hydrography of Kongsfjorden, Svalbard. J. Geophys. Res. Ocean. 110: 1–18. doi:10.1029/2004JC002757

Davis, N. M., D. M. Proctor, S. P. Holmes, D. A. Relman, and B. J. Callahan. 2018. Simple statistical identification and removal of contaminant sequences in marker-gene and metagenomics data. Microbiome 6: 226. doi:10.1186/s40168-018-0605-2

Deng, L., S. Cheung, Z. Xu, K. Liu, and H. Liu. 2023. Microzooplankton Grazing Exerts a Strong Top-Down Control on Unicellular Cyanobacterial Diazotrophs. J. Geophys. Res. Biogeosciences 128: e2023JG007824. 10.1029/2023JG007824

Edler, D., J. Klein, A. Antonelli, and D. Silvestro. 2021. raxmlGUI 2.0: A graphical interface and toolkit for phylogenetic analyses using RAxML. Methods Ecol. Evol. 12: 373–377. 10.1111/2041-210X.13512

Farnelid, H., A. F. Andersson, S. Bertilsson, and others. 2011. Nitrogenase Gene Amplicons from Global Marine Surface Waters Are Dominated by Genes of Non-Cyanobacteria. PLoS One 6: e19223. doi:10.1371/journal.pone.0019223

Farnelid, H., K. Turk-Kubo, M. Muñoz-Marín, and J. Zehr. 2016. New insights into the ecology of the globally significant uncultured nitrogen-fixing symbiont UCYN-A. Aquat. Microb. Ecol. 77: 125–138. doi:10.3354/ame01794

Fernández-Méndez, M., K. A. Turk-Kubo, P. L. Buttigieg, J. Z. Rapp, T. Krumpen, J. P. Zehr, and A. Boetius. 2016. Diazotroph Diversity in the Sea Ice, Melt Ponds, and Surface Waters of the Eurasian Basin of the Central Arctic Ocean. Front. Microbiol. 7: 1–18. doi:10.3389/fmicb.2016.01884

Frank, I. E., K. A. Turk-Kubo, and J. P. Zehr. 2016. Rapid annotation of *nifH* gene sequences using classification and regression trees facilitates environmental functional gene analysis. Environ. Microbiol. Rep. 8: 905–916. doi:10.1111/1758-2229.12455

Frias-Lopez, J., A. Thompson, J. Waldbauer, and S. W. Chisholm. 2009. Use of stable isotope-labelled cells to identify active grazers of picocyanobacteria in ocean surface waters. Environ. Microbiol. 11: 512–525. doi:10.1111/j.1462-2920.2008.01793.x

von Friesen, L. W., M. L. Paulsen, O. Müller, F. Gründger, and L. Riemann. 2023. Glacial meltwater and seasonality influence community composition of diazotrophs in Arctic coastal and open waters. FEMS Microbiol. Ecol. 99: 1–14. doi:10.1093/femsec/fiad067

von Friesen, L. W., and L. Riemann. 2020. Nitrogen Fixation in a Changing Arctic Ocean: An Overlooked Source of Nitrogen? Front. Microbiol. 11. doi:10.3389/fmicb.2020.596426

Gärtner, A., J. Wiese, and J. F. Imhoff. 2008. *Amphritea atlantica* gen. nov., sp. nov., a gammaproteobacterium from the Logatchev hydrothermal vent field. Int. J. Syst. Evol. Microbiol. 58: 34–39. 10.1099/ijs.0.65234-0

Gasol, J. M., and X. A. G. Morán. 2015. Flow Cytometric Determination of Microbial Abundances and Its Use to Obtain Indices of Community Structure and Relative Activity, p. 159–187. *In* T.J. McGenity, K.N. Timmis, and B. Nogales [eds.]. Springer Berlin Heidelberg.

Ghiglione, J. F., P. E. Galand, T. Pommier, and others. 2012. Pole-to-pole biogeography of surface and deep marine bacterial communities. Proc. Natl. Acad. Sci. U. S. A. 109: 17633–17638. doi:10.1073/pnas.1208160109

Gloor, G. B., J. M. Macklaim, V. Pawlowsky-Glahn, and J. J. Egozcue. 2017. Microbiome Datasets Are Compositional: And This Is Not Optional. Front. Microbiol. 8. doi:10.3389/fmicb.2017.02224

Gradoville, M. R., D. Bombar, B. C. Crump, R. M. Letelier, J. P. Zehr, and A. E. White. 2017. Diversity and activity of nitrogen-fixing communities across ocean basins. Limnol. Oceanogr. 62: 1895–1909. doi:10.1002/lno.10542

Gradoville, M. R., H. Farnelid, A. E. White, and others. 2020. Latitudinal constraints on the abundance and activity of the cyanobacterium UCYN-A and other marine diazotrophs in the North Pacific. Limnol. Oceanogr. 65: 1858–1875. doi:10.1002/lno.11423

Halm, H., P. Lam, T. G. Ferdelman, G. Lavik, T. Dittmar, J. LaRoche, S. D’hondt, and M. M. M. Kuypers. 2012. Heterotrophic organisms dominate nitrogen fixation in the South Pacific Gyre. ISME J. 6: 1238–1249.

Harding, K., K. A. Turk-Kubo, R. E. Sipler, M. M. Mills, D. A. Bronk, and J. P. Zehr. 2018. Symbiotic unicellular cyanobacteria fix nitrogen in the Arctic Ocean. Proc. Natl. Acad. Sci. 115: 13371–13375. doi:10.1073/pnas.1813658115

Karl, D., R. Letelier, L. Tupas, J. Dore, J. Christian, and D. Hebel. 1997. The role of nitrogen fixation in biogeochemical cycling in the subtropical North Pacific Ocean. Nature 388: 533–538. doi:10.1038/41474

Karl, D., A. Michaels, B. Bergman, and D. Capone. 2002. The Nitrogen Cycle at Regional to Global Scales, E.W. Boyer and R.W. Howarth [eds.]. Springer Netherlands.

Knapp, A. N. 2012. The sensitivity of marine N_2_ fixation to dissolved inorganic nitrogen. Front. Microbiol. 3. doi:10.3389/fmicb.2012.00374

Knapp, A. N., K. L. Casciotti, W. M. Berelson, M. G. Prokopenko, and D. G. Capone. 2016. Low rates of nitrogen fixation in eastern tropical South Pacific surface waters. Proc. Natl. Acad. Sci. 113: 4398–4403. doi:10.1073/pnas.1515641113

Lehtimaki, J., P. Moisander, K. Sivonen, and K. Kononen. 1997. Growth, nitrogen fixation, and nodularin production by two Baltic Sea cyanobacteria. Appl. Environ. Microbiol. 63: 1647–1656.

Letunic, I., and P. Bork. 2021. Interactive Tree Of Life (iTOL) v5: an online tool for phylogenetic tree display and annotation. Nucleic Acids Res. 49: W293–W296. doi:10.1093/nar/gkab301

Lewis, K. M., G. L. van Dijken, and K. R. Arrigo. 2020. Changes in phytoplankton concentration now drive increased Arctic Ocean primary production. Science (80-.). 369: 198–202. doi:10.1126/science.aay8380

Marie, D., F. Rigaut-Jalabert, and D. Vaulot. 2014. An improved protocol for flow cytometry analysis of phytoplankton cultures and natural samples. Cytom. Part A 85: 962–968.

McMurdie, P. J., and S. Holmes. 2013. phyloseq: An R Package for Reproducible Interactive Analysis and Graphics of Microbiome Census Data M. Watson [ed.]. PLoS One 8: e61217. doi:10.1371/journal.pone.0061217

Mise, K., Y. Masuda, K. Senoo, and H. Itoh. 2021. Undervalued Pseudo-*nifH* Sequences in Public Databases Distort Metagenomic Insights into Biological Nitrogen Fixers S.G. Tringe [ed.]. mSphere 6. doi:10.1128/msphere.00785-21

Moisander, P. H., R. A. Beinart, M. Voss, and J. P. Zehr. 2008. Diversity and abundance of diazotrophic microorganisms in the South China Sea during intermonsoon. ISME J. 2: 954–967.

Moynihan, M. A. 2020. nifHdada2 GitHub repository. doi:10.5281/ZENODO.4283278

Oksanen, J., G. L. Simpson, F. G. Blanchet, and others. 2022. vegan: Community Ecology Package.

Parada, A. E., D. M. Needham, and J. A. Fuhrman. 2016. Every base matters: Assessing small subunit rRNA primers for marine microbiomes with mock communities, time series and global field samples. Environ. Microbiol. 18: 1403–1414. doi:10.1111/1462-2920.13023

Pierella Karlusich, J. J., E. Pelletier, F. Lombard, and others. 2021. Global distribution patterns of marine nitrogen-fixers by imaging and molecular methods. Nat. Commun. 12: 4160. doi:10.1038/s41467-021-24299-y

Priest, T., W.-J. von Appen, E. Oldenburg, and others. 2023. Atlantic water influx and sea-ice cover drive taxonomic and functional shifts in Arctic marine bacterial communities. ISME J. 17: 1612–1625. doi:10.1038/s41396-023-01461-6

Quast, C., E. Pruesse, P. Yilmaz, J. Gerken, T. Schweer, P. Yarza, J. Peplies, and F. O. Glöckner. 2013. The SILVA ribosomal RNA gene database project: improved data processing and web-based tools. Nucleic Acids Res. 41: D590–6. doi:10.1093/nar/gks1219

Riemann, L., E. Rahav, U. Passow, and others. 2022. Planktonic Aggregates as Hotspots for Heterotrophic Diazotrophy: The Plot Thickens. Front. Microbiol. 13. doi:10.3389/fmicb.2022.875050

Robicheau, B. M., J. Tolman, D. Desai, and J. LaRoche. 2023a. Microevolutionary patterns in ecotypes of the symbiotic cyanobacterium UCYN-A revealed from a Northwest Atlantic coastal time series. Sci. Adv. 9: eadh9768. doi:10.1126/sciadv.adh9768

Robicheau, B. M., J. Tolman, S. Rose, D. Desai, and J. LaRoche. 2023b. Marine nitrogen-fixers in the Canadian Arctic Gateway are dominated by biogeographically distinct noncyanobacterial communities. FEMS Microbiol. Ecol. 99: 1–19. doi:10.1093/femsec/fiad122

Rudels, B., R. Meyer, E. Fahrbach, and others. 2000. Water mass distribution in Fram Strait and over the Yermak Plateau in summer 1997. Ann. Geophys. 18: 687–705. doi:10.1007/s00585-000-0687-5

Salazar, G., L. Paoli, A. Alberti, and others. 2019. Gene expression changes and community turnover differentially shape the global ocean metatranscriptome. Cell 179: 1068–1083. doi:10.1016/j.cell.2019.10.014

Salinero, K. K., K. Keller, W. S. Feil, H. Feil, S. Trong, G. Di Bartolo, and A. Lapidus. 2009. Metabolic analysis of the soil microbe *Dechloromonas aromatica* str. RCB: Indications of a surprisingly complex life-style and cryptic anaerobic pathways for aromatic degradation. BMC Genomics 10: 1–23. doi:10.1186/1471-2164-10-351

Selden, C. R., S. V. Einarsson, K. E. Lowry, K. E. Crider, R. S. Pickart, P. Lin, C. J. Ashjian, and P. D. Chappell. 2022. Coastal upwelling enhances abundance of a symbiotic diazotroph (UCYN-A) and its haptophyte host in the Arctic Ocean. Front. Mar. Sci. 9: 1–8. doi:10.3389/fmars.2022.877562

Shao, Z., and Y. W. Luo. 2022. Controlling factors on the global distribution of a representative marine non-cyanobacterial diazotroph phylotype (Gamma A). Biogeosciences 19: 2939–2952. doi:10.5194/bg-19-2939-2022

Shiozaki, T., D. Bombar, L. Riemann, and others. 2017. Basin scale variability of active diazotrophs and nitrogen fixation in the North Pacific, from the tropics to the subarctic Bering Sea. Global Biogeochem. Cycles 31: 996–1009. doi:10.1002/2017GB005681

Shiozaki, T., A. Fujiwara, M. Ijichi, N. Harada, S. Nishino, S. Nishi, T. Nagata, and K. Hamasaki. 2018. Diazotroph community structure and the role of nitrogen fixation in the nitrogen cycle in the Chukchi Sea (western Arctic Ocean). Limnol. Oceanogr. 63: 2191–2205. doi:10.1002/lno.10933

Shiozaki, T., Y. Nishimura, S. Yoshizawa, H. Takami, K. Hamasaki, A. Fujiwara, S. Nishino, and N. Harada. 2023. Distribution and survival strategies of endemic and cosmopolitan diazotrophs in the Arctic Ocean. ISME J. 17: 1340–1350. doi:10.1038/s41396-023-01424-x

Straub, D., N. Blackwell, A. Langarica-Fuentes, A. Peltzer, S. Nahnsen, and S. Kleindienst. 2020. Interpretations of Environmental Microbial Community Studies Are Biased by the Selected 16S rRNA (Gene) Amplicon Sequencing Pipeline. Front. Microbiol. 11.

Swedish Polar Research Secretariat. 2022. SAS 2021 - Meteorological, Oceanographic and Ship Data Collected Onboard Icebreaker Oden. doi:10.48515/0v1w-8958

Thiele, J., and B. Markussen. 2012. Potential of GLMM in modelling invasive spread. CAB Rev. Perspect. Agric. Vet. Sci. Nutr. Nat. Resour. 7: 1–10. doi:10.1079/PAVSNNR20127016

Thompson, A., B. J. Carter, K. Turk-Kubo, F. Malfatti, F. Azam, and J. P. Zehr. 2014. Genetic diversity of the unicellular nitrogen-fixing cyanobacteria UCYN-A and its prymnesiophyte host. Environ. Microbiol. 16: 3238–3249. doi:10.1111/1462-2920.12490

Tschitschko, B., M. Esti, M. Philippi, and others. 2024. Rhizobia–diatom symbiosis fixes missing nitrogen in the ocean. Nature 630: 899–904. doi:10.1038/s41586-024-07495-w

Turk-Kubo, K. A., M. R. Gradoville, S. Cheung, F. M. Cornejo-Castillo, K. J. Harding, M. Morando, M. Mills, and J. P. Zehr. 2022. Non-cyanobacterial diazotrophs: global diversity, distribution, ecophysiology, and activity in marine waters. FEMS Microbiol. Rev. 1–25. doi:10.1093/femsre/fuac046

Turk-Kubo, K. A., H. M. Farnelid, I. N. Shilova, B. Henke, and J. P. Zehr. 2017. Distinct ecological niches of marine symbiotic N_2_-fixing cyanobacterium Candidatus *Atelocyanobacterium thalassa* sublineages. J. Phycol. 53: 451–461. doi:10.1111/jpy.12505

Vihtakari, M. 2022. ggOceanMaps: Plot Data on Oceanographic Maps using “ggplot2.”

Wei, T., and V. Simko. 2021. R package “corrplot”: Visualization of a Correlation Matrix.

Wickham, H. 2016. ggplot2: Elegant Graphics for Data Analysis. Springer-Verlag New York.

Zani, S., M. T. Mellon, J. L. Collier, and J. P. Zehr. 2000. Expression of *nifH* genes in natural microbial assemblages in Lake George, New York, detected by reverse transcriptase PCR. Appl. Environ. Microbiol. 66: 3119–3124. doi:10.1128/AEM.66.7.3119-3124.2000

Zehr, J. P., and D. G. Capone. 2020. Changing perspectives in marine nitrogen fixation. Science (80-.). 368: eaay9514. doi:10.1126/science.aay9514

Zehr, J. P., and L. A. McReynolds. 1989. Use of degenerate oligonucleotides for amplification of the *nifH* gene from the marine cyanobacterium *Trichodesmium thiebautii*. Appl. Environ. Microbiol. 55: 2522–2526. doi:10.1128/aem.55.10.2522-2526.1989

Zehr, J. P., and P. J. Turner. 2001. Nitrogen fixation: Nitrogenase genes and gene expression, p. 271–286. *In* Methods in Microbiology.

