## Supporting Information for "From temperate to polar waters: Transition to non-cyanobacterial diazotrophy upon entering the Atlantic gateway of the Arctic Ocean"

**Affiliations**

**Table S1. Sampling information for each station.** See Materials and Methods in the main text for the rationale of time categories. The coordinates are presented in decimal degrees. AW: Atlantic Water, ArW: Arctic Water, CI: Coastal Influenced Water, UTC: coordinated universal time.

| Station | Latitude (N) | Longitude (E) | Date | Season | Sampling time (UTC) | Time category | Province |
|---|---|---|---|---|---|---|---|
| TR01 | 71.250 | 7.674 | 29/07/2021 | Summer | 17.26 | Afternoon | AW |
| TR02 | 71.825 | 7.825 | 29/07/2021 | Summer | 20.25 | Evening | AW |
| TR03 | 73.380 | 8.245 | 30/07/2021 | Summer | 05.06 | Morning | AW |
| TR04 | 74.708 | 8.533 | 30/07/2021 | Summer | 13.00 | Afternoon | AW |
| TR05 | 76.087 | 8.922 | 30/07/2021 | Summer | 20.56 | Evening | AW |
| TR06 | 77.526 | 8.979 | 31/07/2021 | Summer | 05.06 | Morning | AW |
| TR07 | 78.956 | 8.932 | 31/07/2021 | Summer | 13.05 | Afternoon | ArW |
| TR08 | 80.352 | 9.165 | 31/07/2021 | Summer | 20.53 | Evening | ArW |
| TR09 | 81.005 | 14.917 | 01/08/2021 | Summer | 05.02 | Morning | ArW |
| TR11 | 81.230 | 18.501 | 01/08/2021 | Summer | 18.42 | Evening | ArW |
| TR12 | 81.507 | 26.591 | 02/08/2021 | Summer | 08.21 | Morning | ArW |
| TR13 | 81.508 | 26.807 | 02/08/2021 | Summer | 17.05 | Control | ArW |
| TR14 | 81.507 | 26.748 | 02/08/2021 | Summer | 15.17 | Control (CTD) | ArW |
| TR15 | 82.378 | 8.613 | 11/09/2021 | Autumn | 08.20 | Control (CTD) | ArW |
| TR16 | 82.378 | 8.613 | 11/09/2021 | Autumn | 08.15 | Control | ArW |
| TR17 | 79.426 | 7.952 | 12/09/2021 | Autumn | 13.53 | Mid-day | ArW |
| TR18 | 78.474 | 7.527 | 12/09/2021 | Autumn | 19.38 | Sunset | AW |
| TR19 | 77.028 | 7.310 | 13/09/2021 | Autumn | 04.28 | Sunrise | AW |
| TR20 | 75.856 | 8.087 | 13/09/2021 | Autumn | 11.41 | Mid-day | AW |
| TR21 | 74.688 | 10.846 | 13/09/2021 | Autumn | 18.38 | Sunset | AW |
| TR22 | 73.031 | 14.882 | 14/09/2021 | Autumn | 04.31 | Sunrise | AW |
| TR23 | 71.777 | 16.411 | 14/09/2021 | Autumn | 11.29 | Mid-day | CI |
| TR24 | 70.279 | 15.294 | 14/09/2021 | Autumn | 18.46 | Sunset | CI |
| TR25 | 68.866 | 11.710 | 15/09/2021 | Autumn | 04.19 | Sunrise | CI |
| TR26 | 68.020 | 9.672 | 15/09/2021 | Autumn | 10.10 | Mid-day | CI |
| TR27 | 66.738 | 7.971 | 15/09/2021 | Autumn | 17.50 | Sunset | CI |
| TR28 | 64.765 | 5.883 | 16/09/2021 | Autumn | 04.40 | Sunrise | CI |
| TR29 | 63.366 | 4.409 | 16/09/2021 | Autumn | 12.16 | Mid-day | CI |
| TR30 | 62.186 | 4.003 | 16/09/2021 | Autumn | 18.13 | Sunset | CI |
| TR31 | 60.017 | 4.001 | 17/09/2021 | Autumn | 05.00 | Sunrise | CI |
| TR32 | 59.211 | 4.022 | 17/09/2021 | Autumn | 11.30 | Mid-day | CI |
| TR33 | 58.225 | 5.481 | 17/09/2021 | Autumn | 17.53 | Sunset | CI |
| TR34 | 57.703 | 8.252 | 18/09/2021 | Autumn | 05.04 | Sunrise | CI |
| TR35 | 57.808 | 9.901 | 18/09/2021 | Autumn | 10.54 | Mid-day | CI |
| TR36 | 57.517 | 11.330 | 18/09/2021 | Autumn | 16.55 | Sunset | CI |

**Table S2. Primers for amplicon sequencing.** PCR primers applied in this study for the amplification of *nifH* (nifH1-4) and 16S rRNA (515F-Y, 926R) genes (Zehr and McReynolds 1989; Zani et al. 2000; Parada et al. 2016).

| Primer name | Sequence (5'-3') | Reference |
|---|---|---|
| nifH1 | TGYGAYCCNAARGCNGA | Zehr & McReynolds, 1989 |
| nifH2 | ADNGCCATCATYTCNCC | |
| nifH3 | ATRTTRTTNGCNGCRTA | Zani et al., 2000 |
| nifH4 | TTYTAYGGNAARGGNGG | |
| 515F-Y | GTGYCAGCMGCCGCGGTAA | Parada et al., 2016 |
| 926R | CCGYCAATTYMTTTRAGTTT | |

**Table S3. Primers and probes for qPCR.** Primer/probe sets applied in this study for quantitative PCR (qPCR) and reverse-transcription qPCR (RT-qPCR) of three non-cyanobacterial diazotroph groups (Gamma-Arctic1, Gamma-Arctic2, and Beta-Arctic1; developed in this study, see "qPCR and RT-qPCR of *nifH*" in the main text) and three previously designed assays (UCYN-A1, UCYN-A2/A4, and *Nodularia* sp.) (Church et al. 2005; Boström et al. 2007; Thompson et al. 2014). $R^2$ is the standard curve correlation coefficient of determination and E is the efficiency (%) of the assay. The Limit of Detection (LOD) and Limit of Quantification (LOQ) are presented as *nifH* gene/transcript copies $L^{-1}$ of seawater.

| Target | Forward primer (5'-3') | Reverse primer (5'-3') | Probe (5'-3') | Targeted ASVs (top 50) | Reference | $R^2$ | E (%) | LOD (copies $L^{-1}$) | LOQ (copies $L^{-1}$) |
|---|---|---|---|---|---|---|---|---|---|
| UCYN-A1 | GGCTATAACAACGTTTTATGCGTTGA | ACCACGACCAGCACATCCA | TCCGGTGGTCCTGAGCCTGGA | 3, 27, 39, 41, 49, 50, 51 | Church et al. 2005 | 0.994 | 86 | 122 ± 69 | 146 ± 95 |
| UCYN-A2/A4 | GGTTACAACAACGTTTTATGTGTTGA | ACCACGACCAGCACATCCA | TCTGGTGGTCCTGAGCCCGGA | 4, 33, 37 | Thompson et al. 2014 | 0.995 | 87 | 170 ± 145 | 193 ± 164 |
| *Nodularia* sp. | CGAAGAAGTAATGCTGACCGG | CGGATAGGCATAGCAAAACCA | ACCCGGTGTAGGTTGTGCTGGTCG | 9 | Boström et al. 2007 | 0.998 | 86 | 164 ± 140 | 186 ± 152 |
| Beta-Arctic1 | ATCAYCGCCATCAACTTCC | ACGATGTAGATTTCCTGAGCC | CGGTGGCTTCGCCATGCCGAT | 5, 12, 16, 48 | This study | 0.984 | 114 | 137 ± 119 | 141 ± 123 |
| Gamma-Arctic1 | ATTATGGAAATGGCGGCAG | GACCACCAGATTCAACACAG | CGGCACCGTAGAAGACCTTGAACTTGAA | 1, 2, 19, 21, 26, 31, 32, 43, 45, 46 | This study | 0.991 | 109 | 85 ± 48 | 114 ± 64 |
| Gamma-Arctic2 | ATCATGGAGATGGCTG | GACCACCGGACTCAACACAC | AGGTACTGTAGAAGATCTGGAACTGGAA | 6 | This study | 0.998 | 94 | 325 ± 133 | 336 ± 138 |

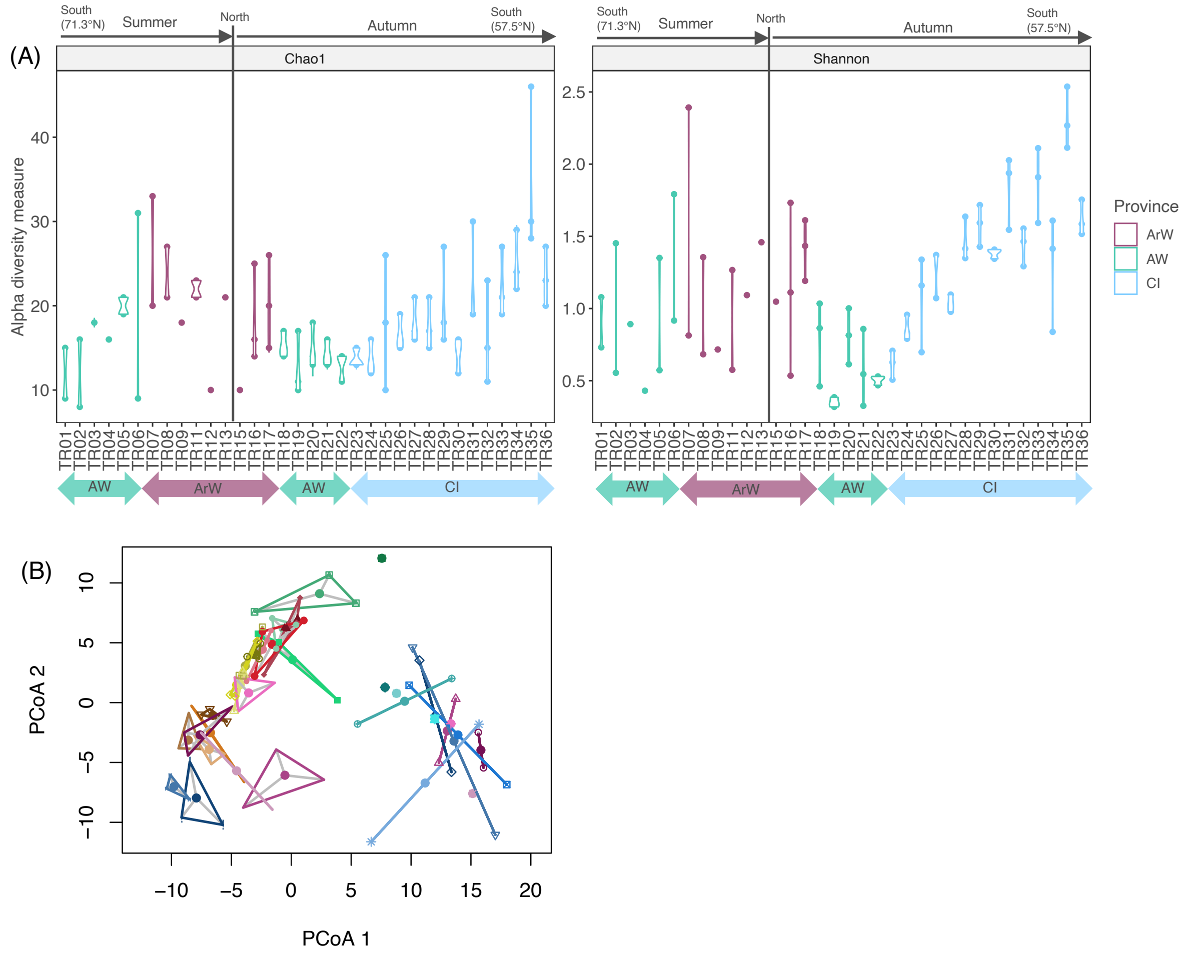


**Fig. S1. Variation in *nifH* amplicon sequence variant (ASV) composition between replicate samples from each station. (A)** Violin plots of Chao1 and Shannon alpha-diversity indices for each station (one to three replicates per station). ArW: Arctic Water, AW: Atlantic Water, CI: Coastal Influenced Water. **(B)** Principal Coordinate analysis (PCoA) ordination displaying the two first components (PCoA1 and PCoA2) of multivariate homogeneity of group dispersion (betadisper (); Aitchison distance). Each colored dot represents the center of variation for a station and the same-colored lines (if two replicates) or triangles (if three replicates) the variance among replicates.

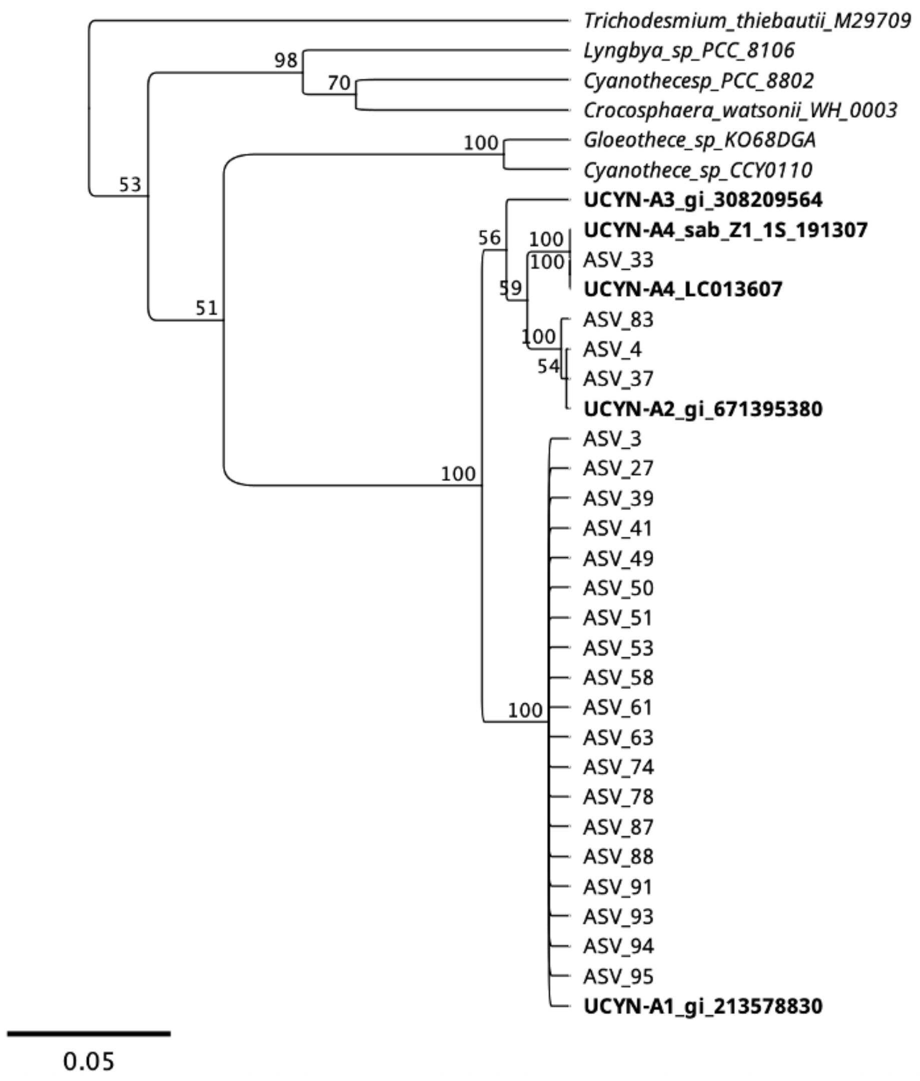


**Fig. S2. UCYN-A sublineage affiliation.** UPGMA (unweighted pair group method with arithmetic mean)-nucleotide tree of all Chroococcales-assigned ASVs (from the top 100 ASVs) with UCYN-A sublineage reference sequences in bold (Farnelid et al. 2016; Turk-Kubo et al. 2017). Based on Jukes-Cantor distances, geneious alignment, and 1000 bootstraps (presented as values from 0-100). Performed in Geneious Prime (v.1.1).

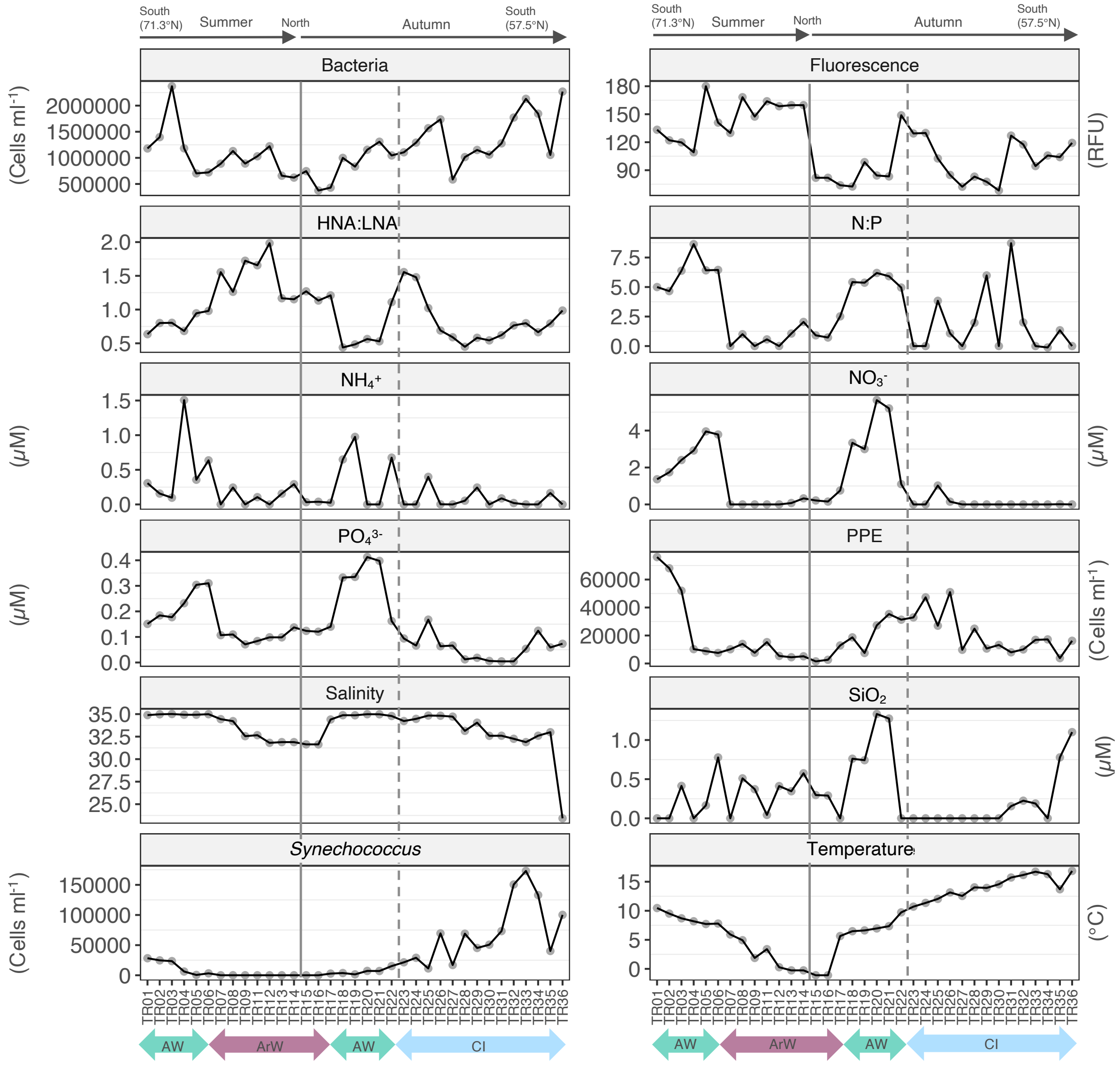


**Fig. S3. Environmental conditions along the summer and autumn transects.** HNA:LNA is the ratio of High Nucleic Acid to Low Nucleic Acid bacterial cells (proxy of bacterial activity), N:P is the molar ratio between dissolved inorganic nitrogen (nitrate, nitrite, ammonium) and dissolved inorganic phosphorous (phosphate). $NH_4^+$: ammonium, $NO_3^-$: nitrate + nitrite, $PO_4^{3-}$: phosphate, PPE: photosynthetic picoeukaryotes, $SiO_2$: silicate. The dashed line indicates the transition from province AW to CI (station TR22 to TR23) and the solid line indicates the division between the summer and autumn transects.

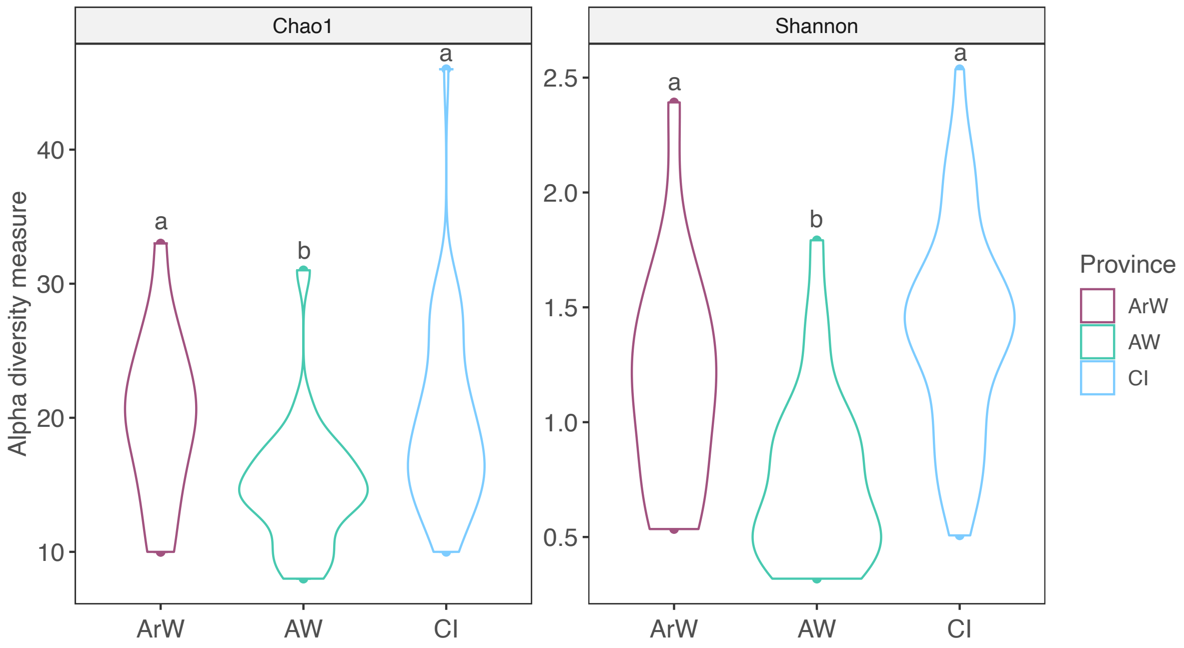


**Fig. S4. Alpha diversity indices of *nifH* amplicon sequence variants in the different provinces.** Violin plots of the average Chao1 and Shannon alpha diversity indices in the three provinces (summer and autumn transects merged). Different letters denote statistical significance ($p<0.05$). ArW: Arctic Water, AW: Atlantic Water, CI: Coastal Influenced Water.

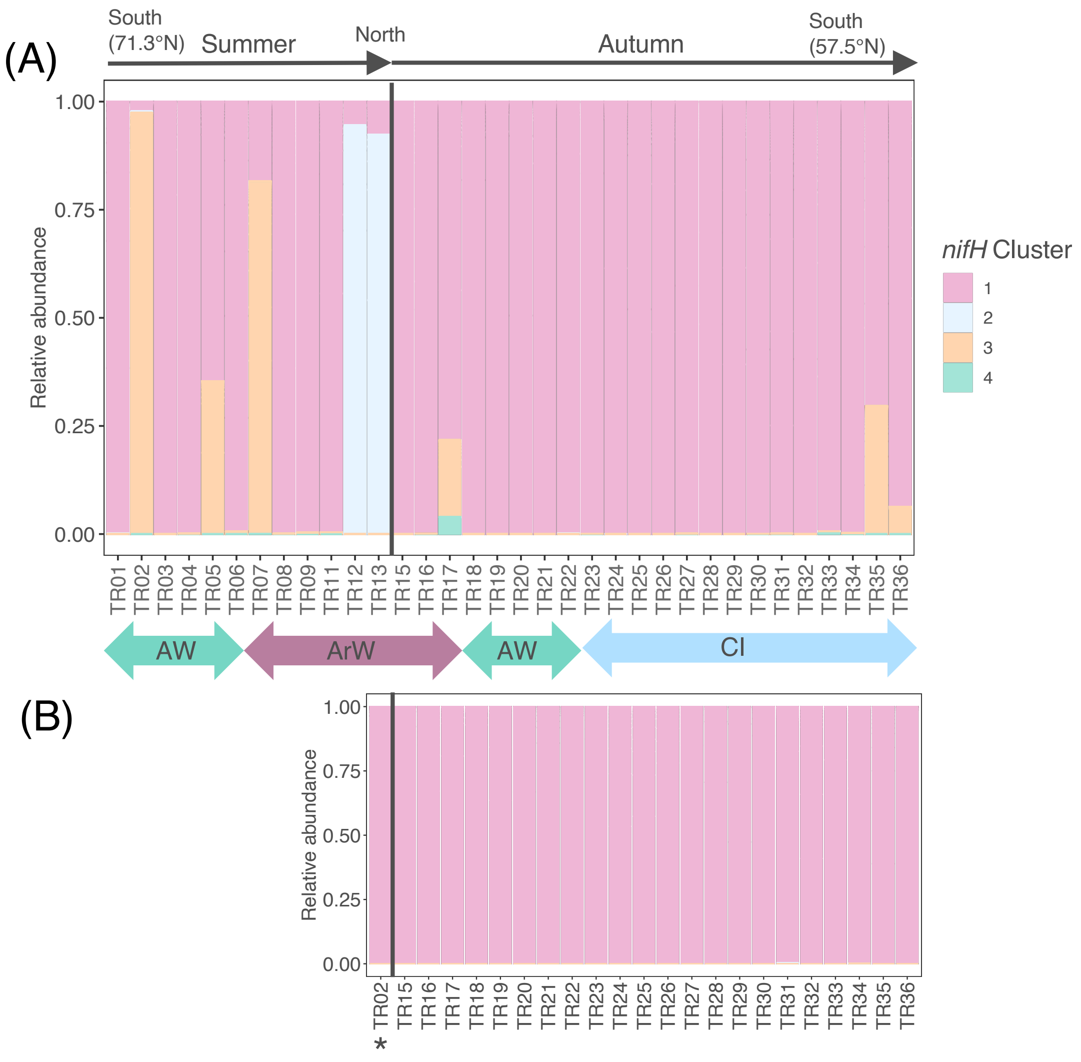


**Fig. S5. Relative abundance of *nifH* clusters. (A)** DNA (present *nifH*) along the summer and autumn transects (one to three replicates merged for each station), and **(B)** cDNA (transcribed *nifH*) along the autumn transect. The asterisk in B denotes station TR02 from summer AW (the only successful *nifH* amplification from cDNA in summer). ArW: Arctic Water, AW: Atlantic Water, CI: Coastal Influenced Water.

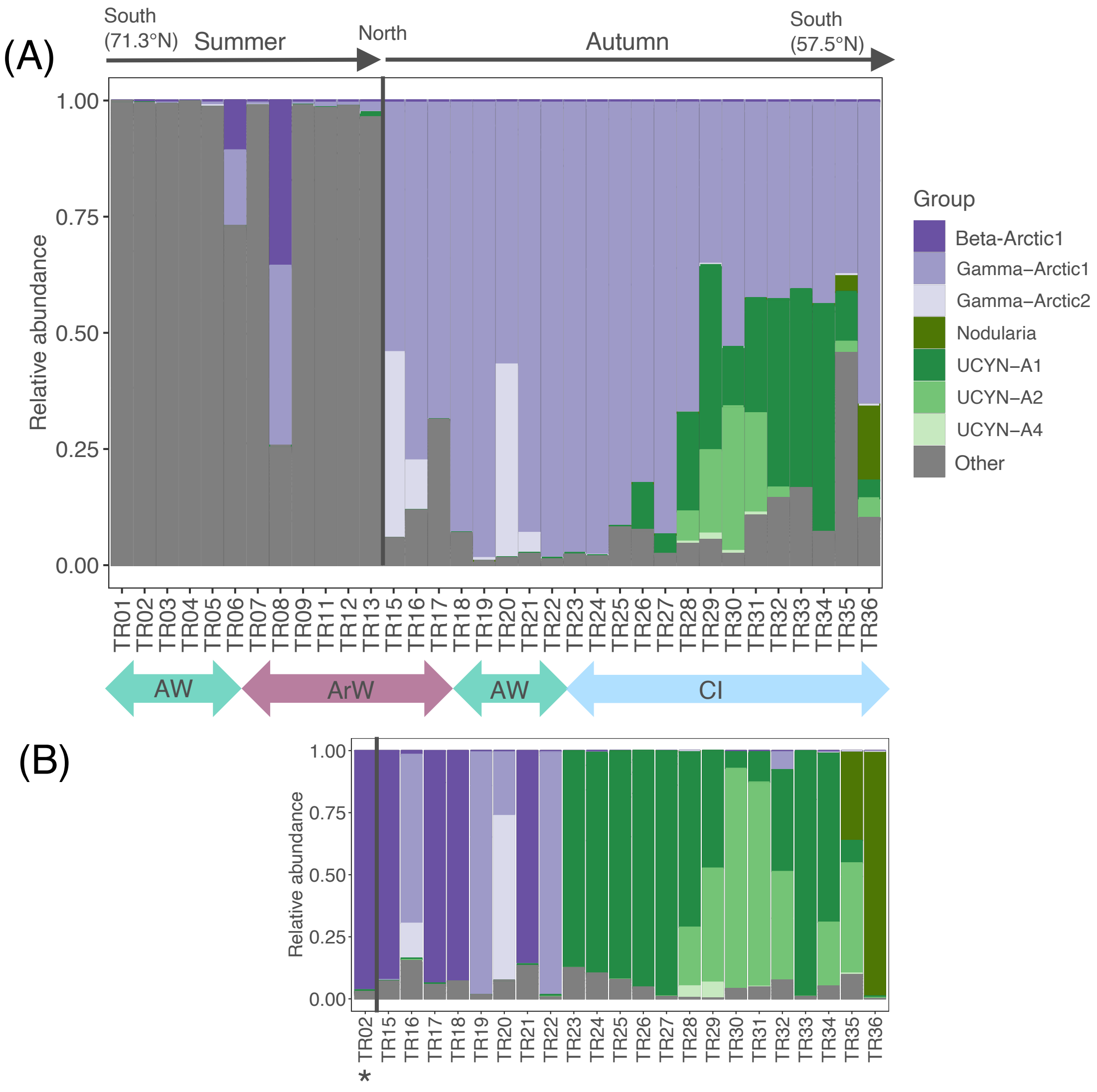


**Fig. S6. Relative abundance of UCYN-A1, UCYN-A2, UCYN-A4, *Nodularia* sp., Gamma-Arctic1, Gamma-Arctic2 and Beta-Arctic1** (*nifH* amplicon sequencing). **(A)** DNA (present *nifH*) along the summer and autumn transects (one to three replicates merged for each station), and **(B)** cDNA (transcribed *nifH*) along the autumn transect. The asterisk denotes sample TR02 from summer AW (the only successful *nifH* amplification from cDNA in summer). ArW: Arctic Water, AW: Atlantic Water, CI: Coastal Influenced Water, NCD: non-cyanobacterial diazotrophs.

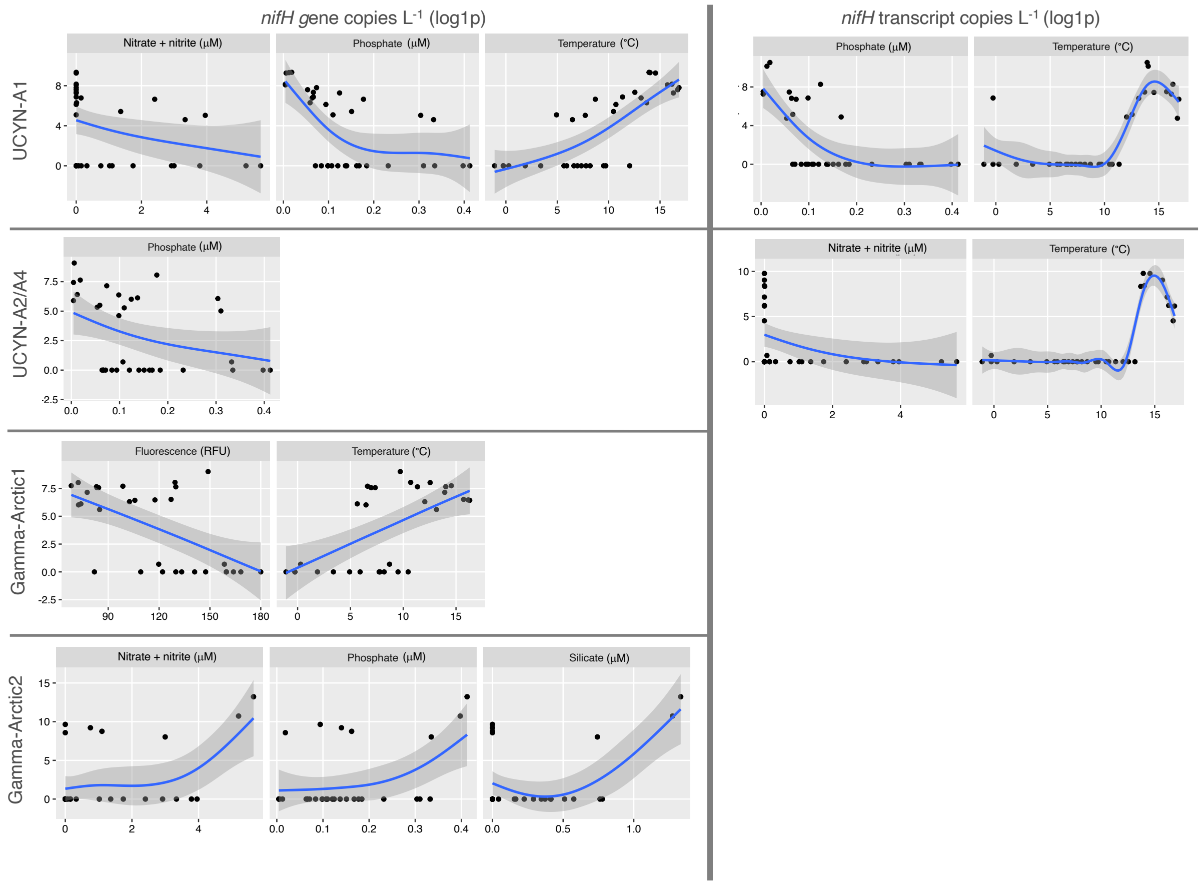


**Fig. S7**. **Correlation of diazotrophs with environmental variables.** Significant correlations of diazotroph group-specific *nifH* gene/transcript copies (y-axis) with environmental variables (x-axis) displayed as generalized additive models (GAM) of log1p-transformed (natural logarithm of x+1) *nifH* gene or transcript copies $L^{-1}$ against environmental variables (shading denotes 95% confidence interval around the GAM smoothing). Temperature was not significantly correlated with UCYN-A1 transcript copies but is included here for visualization of the non-monotonic relationship indicating a temperature optimum at ~14°C.

### References

Boström, K. H., L. Riemann, U. L. Zweifel, and Å. Hagström. 2007. *Nodularia* sp. *nifH* gene transcripts in the Baltic Sea proper. J. Plankton Res. 29: 391–399. doi:10.1093/plankt/fbm019

Church, M. J., B. D. Jenkins, D. M. Karl, and J. P. Zehr. 2005. Vertical distributions of nitrogen-fixing phylotypes at Stn ALOHA in the oligotrophic North Pacific Ocean. Aquat. Microb. Ecol. doi:10.3354/ame038003

Farnelid, H., K. Turk-Kubo, M. Muñoz-Marín, and J. Zehr. 2016. New insights into the ecology of the globally significant uncultured nitrogen-fixing symbiont UCYN-A. Aquat. Microb. Ecol. 77: 125–138. doi:10.3354/ame01794

Parada, A. E., D. M. Needham, and J. A. Fuhrman. 2016. Every base matters: assessing small subunit rRNA primers for marine microbiomes with mock communities, time series and global field samples. Environ. Microbiol. 18: 1403–1414. doi:10.1111/1462-2920.13023

Thompson, A., B. J. Carter, K. Turk-Kubo, F. Malfatti, F. Azam, and J. P. Zehr. 2014. Genetic diversity of the unicellular nitrogen-fixing cyanobacteria UCYN-A and its prymnesiophyte host. Environ. Microbiol. 16: 3238–3249. doi:10.1111/1462-2920.12490

Turk-Kubo, K. A., H. M. Farnelid, I. N. Shilova, B. Henke, and J. P. Zehr. 2017. Distinct ecological niches of marine symbiotic $N_2$-fixing cyanobacterium Candidatus *Atelocyanobacterium thalassa* sublineages. J. Phycol. 53: 451–461. doi:10.1111/jpy.12505

Zani, S., M. T. Mellon, J. L. Collier, and J. P. Zehr. 2000. Expression of *nifH* genes in natural microbial assemblages in Lake George, New York, detected by reverse transcriptase PCR. Appl. Environ. Microbiol. 66: 3119–3124. doi:10.1128/AEM.66.7.3119-3124.2000

Zehr, J. P., and L. A. McReynolds. 1989. Use of degenerate oligonucleotides for amplification of the *nifH* gene from the marine cyanobacterium *Trichodesmium thiebautii*. Appl. Environ. Microbiol. 55: 2522–2526. doi:10.1128/aem.55.10.2522-2526.1989
